# An evolutionarily conserved olfactory receptor is required for sex differences in blood pressure

**DOI:** 10.1101/2022.11.16.516677

**Authors:** Jiaojiao Xu, Rira Choi, Kunal Gupta, Helen R. Warren, Lakshmi Santhanam, Jennifer L. Pluznick

## Abstract

Sex differences in blood pressure are well-established, with premenopausal women having lower blood pressure than men by ∼10mmHg; however, the underlying mechanisms are not fully understood. We report here that olfactory receptor 558 (Olfr558), which has not previously been studied in non-olfactory tissues, localizes to vascular smooth muscle cells in numerous tissues including the kidney and heart. In the kidney, Olfr558 colocalizes with renin (a hormone that plays a key role in blood pressure regulation) in the renal afferent arteriole. Based on the localization of Olfr558, we hypothesized that Olfr558 plays a role in blood pressure regulation. We find that sex differences in blood pressure are intact in Olfr558 wildtype (WT) mice, but, are absent in Olfr558 knockout (KO) mice. We find that male KO mice have lowered diastolic blood pressure, decreased renin expression and activity, and altered vascular reactivity. Female KO mice exhibit increased blood pressure and increased pulse wave velocity, indicating increased vascular stiffness. The human ortholog of Olfr558, OR51E1, was previously identified as a locus associated with diastolic blood pressure. We report here that a rare OR51E1 missense variant has a statistically significant sex interaction effect with diastolic blood pressure, increasing diastolic blood pressure in women but decreasing it in men. In addition, we characterize how two different clinically relevant OR51E1 variants influence OR51E1 signaling *in vitro*. In sum, our findings demonstrate an evolutionarily conserved role for Olfr558/OR51E1 to mediate sex differences in blood pressure by altering renin, vascular reactivity, and arterial stiffness.

## Introduction

Sex differences in blood pressure are well-established, with premenopausal women having lower blood pressure than men by ∼10mmHg^1-14^. These sex differences are modeled well in C57BL/6 mice, where females have mean arterial pressure ∼10 mmHg lower than males by telemetry^1^. However, the clinical definition of hypertension is identical in men and women (>130/80 for systolic pressure/ diastolic pressure), in part because the mechanisms underlying sex differences in blood pressure regulation are only partially understood. Although sex hormones likely contribute to sex differences in blood pressure, they are not fully responsible: hormone replacement therapy in postmenopausal women does not consistently alter blood pressure^15-17^, and lowered testosterone in aging men is associated with increased risk of cardiovascular diseases^18-20^. Similarly, sex chromosomes may contribute to sex differences in blood pressure, but cannot fully explain these differences^21^.

Olfactory receptors (ORs), the largest family of G-protein-coupled-receptors (GPCRs), act in the olfactory epithelium to mediate the sense of smell^22^. However, ORs are also expressed and play functional roles in non-olfactory tissues such as kidney, eyes, prostate, and sperm^23-25^. Although there are 1000 murine ORs in the genome, olfactory receptor 558 (Olfr558) is a particularly promising candidate for study. First, Olfr558 is one of only 3 mammalian ORs to have been well-conserved by evolution: Olfr558 has a clear ortholog in all placental mammals, including mice, rats, rabbits, elephants, horses, and 5 species of primates^26-29^, and the human ortholog (olfactory receptor family 51 subfamily E member 1; OR51E1) has one of the lowest rates of single nucleotide polymorphism among human ORs^30^. Second, OR51E1 is widely expressed outside the olfactory epithelium in humans (in 13 different tissues^31^), and we previously reported that Olfr558 is expressed in murine kidney^32,33^. Third, most ORs are orphan receptors with no identified ligands; however, we have previously identified ligands for Olfr558, with the strongest ligand being butyrate, one of the major short-chain fatty acids (SCFAs) produced by gut microbiome^33,34^. Thus, we undertook a study to uncover the physiological role of Olfr558 and found that this evolutionarily conserved receptor has a novel role to mediate sex differences in blood pressure.

## Methods

### Ethical approval

All animal protocols and procedures were approved by the Johns Hopkins University Institutional Animal Care and Use Committee (accredited by the Association for Assessment and Accreditation of Laboratory Animal Care International).

### Animals

Global Olfr558 knockout (KO) mice were generated by Ingenious Targeting Laboratory, using Flexible Accelerated STOP TetO (F.A.S.T) technology^35^ by inserting a stop cassette prior to the ATG start site of Olfr558. Olfr558 heterozygotes (C57BL6/N background) were bred in-house to obtain Olfr558 wild-type (WT) and KO littermates. Mice were housed in individually ventilated cages with maximum of five adult mice per cage. Genotypes were confirmed by PCR of tail biopsies. All animals were given unrestricted access to food and water throughout the duration of the experiments and were maintained on Teklad 2018SX, 18% protein diet. For radio-telemetry studies, mice were housed separately in static cages placed on the telemetry receiver platform.

### qPCR

Olfr558 WT mice were sacrificed between 10 am to 4 pm. After sacrifice, tissues of interest were homogenized in Trizol (Life Technologies) and RNA was extracted using the RNeasy mini kit (Qiagen). Isolated RNA was reverse transcribed to cDNA using reverse transcriptase (RT-PCR) or water (mock RT-PCR). qPCR was performed using Taqman Master Mix (Applied Biosystems) and probes for: Olfr558 (Cat.NO. Mm012798500_m1, Thermo Fisher), renin (Cat. NO. Mm02342887_mH, Thermo Fisher), Akr1b7 (Cat. NO. Mm00477605_m1, Thermo Fisher, renin cell marker), or GAPDH (Cat. NO. Mm99999919_g1, Thermo Fisher) by cycling at 50°C for 2 min, 95 °C 10 min, and followed by 95 °C 15 s and 60 °C 1 min for 40 cycles. RT-PCR and mock RT-PCR were run simultaneously.

### RNAScope and immunostaining

Olfr558 localization was determined by a combination of RNAScope and immunostaining. Olfr558 WT and KO mice under anesthesia were perfused with PBS, followed by 4% paraformaldehyde (PFA) (Cat. No.15714-S, Electron Microscopy Sciences) between 10 am to 4 pm. Tissues including kidney, heart, brown adipose tissue, and skeletal muscle were harvested, fixed in 4% PFA at 4°C overnight, and then immersed in 30% sucrose. For olfactory epithelium (OE), the head (with fur, skin and lower jaw removed) was fixed in 4% PFA at 4°C overnight and then transferred to PBS overnight. The head was immersed in EDTA solution (120 g EDTA in 900 ml H_2_O with pH=7.3) at 4°C for one week, and then dehydrated with 30% sucrose (30 g sucrose in 100 ml PBS), 50%OCT/50% sucrose, and finally 100% OCT for 48 h, 20-30 min, and 20-30 min, respectively.

For all tissues, the tissue was then frozen in OCT and 10 μm cryo-sections were generated. RNAScope was then performed. For RNAScope, 10 μm cryosections were permeabilized, retrieved, and hybridized with Mm-Olfr558 probe (Cat. No. 316131, ACDBio), followed by AMP (RNAScope@ Multiplex FL v2 AMP, ACDBio) hybridization and HRP-C1 (Cat. No. 323104, ACDBio) signal development. For immunostaining, sections were then blocked with superblock, and incubated with rabbit anti-alpha-smooth muscle actin (α-SMA, Cat. NO. ab5694, Abcam) or rabbit anti-renin (a generous gift from Dr. Tadashi Inagami, Vanderbilt) antibody overnight at 4°C. Sections were then washed and incubated with Alexa-fluor 488 goat-anti rabbit secondary antibody followed by Hoechst staining. Sections were imaged using a confocal microscope (Zeiss LSM 700). For renin-glomeruli association, glomeruli were identified by Hoescht staining, then scored (+/-) for adjacent renin signal. Approximately 90% glomeruli in each section were scored. Images were taken using a fluorescence microscope (Keyence BZ-X710).

Olfr558 and Olfr78 probe double staining were examined using RNAScope. Briefly, kidney sections were hybridized with Olfr558 (C1) and Olfr78 (Cat. No. 436601-C2, ACDBio) probes and then signal was developed using HRP-C1 for Olfr558, and subsequently developed using HRP-C2 (Cat. No. 323105, ACDBio) for Olfr78. Images were obtained by a confocal microscope (Zeiss LSM 700).

### Radio-telemetry

Male (3-month old) and female (4-month old) Olfr558 WT and KO mice were implanted with radio-telemetry devices (PA-C10, Data Science International) as previously described ^36,37^. Radio-telemetry catheters were inserted into the aorta from right carotid artery, and the body of transmitter was placed subcutaneously in the abdomen. The procedure was performed under the mixture of 1.5% isoflurane and 2% O_2_. Ten to fourteen days after surgery, telemetry readings began. Blood pressure was measured every 30 min with a duration of 10 s for a total of 5 days after recovery. The light/dark cycle was 7am to 9 pm and 9 pm to 7am, respectively.

### Wire myography

Active properties of the vessels in response to vasoconstrictors and vasodilators were examined by wire myography in both the aorta and mesenteric artery as reported^38^. Briefly, the thoracic aorta was cut into 2-mm rings after carefully removing connective tissues. Each aortic ring was placed into Krebs buffer (containing [in mM] 118 NaCl, 4.7 KCl, 2.5 CaCl_2_, 1.1 KH_2_PO_4_, 25 NaHCO_3_, 1.2 MgSO_4_, and 11 glucose), and then mounted to a myograph chamber (DMT) and continuously bubbled with 95% O2 and 5% CO2 at 37°C. The rings were stretched to a final tension of 550 mg in 100-mg force increments. The 2^nd^ and 3^rd^ order mesenteric arteries were dissected and cut into 2-mm in HEPES-Tyrode solution (containing [in mM] 137 NaCl, 2.7 KCl, 1.8 CaCl_2_, 1 MgCl_2_·6H_2_O, 5.6 D-glucose, and 10 HEPES, pH 7.3-7.4) as previously reported^39^. Two wires were inserted into the lumen of the mesenteric artery in HEPES-Tyrode solution in a petri-dish, and the vessel segments were mounted on the wire myograph chamber. All mounted vessels were submerged in Krebs buffer. The mesenteric artery was stretched to a final tension of 500 mg in 100-mg force increments. KCl (60 mM for Aorta and 100 mM for mesentery) were added to determine vessel viability and obtain the maximal contractility. Phenylephrine (PE) induced vasoconstriction was studied using increasing doses of PE (10^−9^ to 10^−5^ M). Endothelial-mediated vasorelaxation was then evaluated using increasing doses of acetylcholine (ACh, 10^−9^ to 10^−5^ M) after PE pre-constriction. Finally, endothelial-independent vasorelaxation was measured by increasing doses of sodium nitroprusside (SNP, 10^−9^ to 10^−5^ M) in vessels pre-constricted with PE.

### Pulse wave velocity (PWV)

PWV was measured noninvasively using a high-frequency and high-resolution Doppler spectrum analyzer, as previously reported^38^. Briefly, Olfr558 WT and KO females (10-week-old) were placed supine on a 37°C plate anesthetized with 1.5% isoflurane and 2% O_2_. The aortic pulse wave at thorax and abdomen were separately recorded at a distance of ∼3 cm using a 10 MHz probe, simultaneously with electrocardiograph recording. The time for the pulse wave transit from the thoracic to the abdominal aorta was measured using the electrocardiograph as a fixed point. Subsequently, PWV was calculated as the separation distance divided by the pulse transit time between the two points.

### Tensile testing

The mechanical properties of intact and decellularized aortic rings were measured by tensile testing, as previously reported^40^. Briefly, the thoracic aorta from 10-week-old Olfr558 WT and KO females were harvested and cut into 2-mm rings. Two intact and two decellularized rings from each animal were tested. Vessel dimensions (lumen diameter, wall thickness, and length) were calculated by the transverse and longitudinal images of aortic rings obtained using a light microscope. Each vessel was then mounted on an electromechanical puller (DMT). After calibration, the pins were moved apart using an electromotor, and displacement and force applied by the vessel wall on the pin were continuously recorded. To calculate engineering stress (S), force (F) was normalized to the initial stress-free area of the specimen (S=F/2t × L; t=thickness, L= length of the aortic ring). Engineering strain (λ) was calculated as the ratio of displacement to the initial stress-free diameter. The stress-strain relationship was represented by the equation S=α exp (β λ), where α and β are constants. α and β were determined by nonlinear regression for each sample and used to generate stress-strain curves by treating the x-axis as a continuous variable. Incremental elastic modulus (E_inc_) was calculated as the slope of the stress-strain curve at a strain of 0.5 and 1.8.

### Other *in vivo* studies

Whole blood samples were collected from the superficial temporal vein of 3-month old Olfr558 WT and KO mice, and were analyzed by an iStat Chem8+ cartridge (Abbott). Glomerular filtration rate (GFR) was measured in conscious and unrestrained mice using transcutaneous measurement of FITC-sinistrin (MediBeacon) as previously reported^41,42^. Briefly, a small area on the back of the mouse was shaved and depilated with Nair under anesthesia with a mixture of 1.5% isoflurane and 2% O_2_. A GFR device (MediBeacon), which measures transcutaneous fluorescence, was attached to the hairless region of the back and secured with surgical tape. A 7 mg/100 g body weight of FITC-sinistrin was retro-orbitally injected into the mouse who was then placed back into the cage with free access to water and food for 1.5 h, after which the device was removed and data analyzed using MediBeacon software. Mice body weight on week 3 and 8 were measured, kidney weight/body weight (KW/BW) and heart weight/body weight (HW/BW) were measured as well. All in vivo data were collected from both Olfr558 WT and KO males and females. The “n” for each study is noted in figure legends.

### Plasma renin activity

Plasma renin activity was measured in 3-month old Olfr558 WT and KO mice with a modified angiotensin I measurement kit (S-1188, Peninsula Laboratories). Plasma was collected from male and female Olfr558 WT and KO mice treated with 0.49% NaCl diet (Cat. No. TD.96208 modified with orange color, Envigo). From males, we also collected plasma from mice given Teklad diet (Teklad 2018SX, 18% protein). Plasma was diluted 15-fold, and then incubated with excess porcine angiotensinogen (SCP0021, Sigma) for 20 min at 37°C in a buffer containing 50 mM sodium acetate (pH 6.5), 10 mM 4-(2-aminoethyl)benzenesulfony fluoride hydrochloride (AEBSF, Sigma-Aldrich), 10 mM EDTA (pH 8.0), 1 μM porcine angiotensinogen, and 10 mM 8-hydroxyquinoline (Sigma-Aldrich). After incubation, the sample was analyzed according to the provided protocol. Plasma renin activity was assayed by competitive binding of angiotensin I antibody.

### 16S rRNA Microbiome Analysis

Fecal pellets (1 to 3 for each sample) were collected from 8-week-old Olfr558 WT and KO males and females. Fecal DNA was isolated using the Fast Stool Isolation Kit (Cat. No. 51604, Qiagen). As previously reported^43^16S rRNA V3 through V4 region was amplified using primers 319F (CTCCTACGGGAGGCAGCAGT) and 806R (GGACTACHVGGGTWTCTAAT). Sequencing was performed by the Johns Hopkins Transcriptomics and Deep Sequencing Core and analyzed by Resphera Biosciences as previously^43^. Differential abundance analysis of α-diversity analyzed differences between groups (nonparametric difference test, Mann-Whitney *U* test, and *t* test). Multiple hypothesis testing was corrected using the false discovery rate. Generalized linear modeling was performed using R. 16S data have been deposited to the National Center for Biotechnology Information. (https://www.ncbi.nlm.nih.gov/Traces/study/?acc=PRJNA902003&o=acc_s%3Aa)

### Human Genetic Analysis

A previous genome-wide association study^44^ (GWAS) identified OR51E1 as a locus significantly associated with diastolic blood pressure (DBP). Here, we performed new genetic association analyses of DBP within the UK Biobank (UKB) cohort, in order to conduct more detailed stratified analyses of this published variant, and also to analyze a new rare variant of interest. We used the same dataset that was used for the previous GWAS (Supplementary Table 2), as described in Evangelou et al, 2018. All UKB individuals analyzed are of European ancestry. In order to perform simple linear regression analyses, we excluded 1^st^ and 2^nd^ degree relatives, leaving a total of n=423,657 individuals for analysis. Linear regression analyses testing the association of the genetic variant with DBP were adjusted for the following covariates: sex, age, age^2, BMI, genotyping chip subset of UKB, and top 10 Principal components of ancestry. Sex-stratified analyses were performed using two separate subgroups to compare males vs females. In addition, an analysis was performed on all individuals with a sex-interaction term to identify any sex-specific effects. Finally, given the changes in blood pressure that occur with aging (particularly in women), we also performed age-stratified analyses in women (<50 vs >50 years) and in men (<50 vs >50 years), and a corresponding age-interaction analysis. Within each analyzed subgroup, we calculated the mean DBP measurement for creation of plots. We performed analyses for two variants: (i) the common missense variant, rs17224476 (chr11:4673788, NCBI build 37), previously identified by Evangelou, et al^44^ as the lead variant at the *OR51E1* locus achieving the most significant association for DBP. And, (ii), a rare missense variant, rs202113356, which we previously published to alter OR51E1 function (chr11:4673788, NCBI build 37)^34^. This rare variant had not been analyzed previously by Evangelou et al, as the published GWAS only included common variants. As rs202113356 is a rare variant, there were no individuals in UKB with two copies of the minor allele A. Thus, for consistency, all of our statistical analyses for both variants are performed according to a binary genotype model comparing non-carriers (GG) versus carriers of the minor allele (AG/AA). The common variant (rs17224476) is directly genotyped within the UKB data, whereas the rare variant (rs202113356) is extracted from the imputed genetic data, with high imputation quality Rsq = 0.8. For analysis by genotype groups, the genetic data was converted to hard-call genotype format using PLINK software, yielding N=58 individuals with missing data for the imputed rare variant if their imputation certainty did not reach the hard-call threshold in PLINK.

### Cloning and immunofluorescence

OR51E1 has been previously cloned with Lucy-Flag-Rho tags at the N-terminus in the pME18S vector^45^. Additionally, we previously reported that Lucy- and Rho-tags promote surface expression of olfactory receptors (ORs)^45^. Thus, we used the Lucy-Flag-Rho tags for OR51E1 variants—rs202113356 (OR51E1 A156T) and rs17224476 (OR51E1 S10N). The OR51E1 A156T mutant was generated as part of a previous study^34^; the S10N mutant was generated for this study via PCR, and sequenced to confirm. To examine OR expression and surface trafficking. HEK 293T cells were seeded onto poly-L-lysine coated coverslips and transfected with OR constructs with receptor transporting protein 1S (RTP1S) and/or olfactory G protein (Golf), as we previously reported^46^. Cell surface staining was done in live cells blocked with diluted superblock (1:4 dilution in PBS with 1 mM MgCl_2_ and 0.1 mM CaCl_2_, ThermoFisher Scientific) at 4°C for 0.5 h, and incubated with rabbit polyclonal anti-flag antibody (Cat. No. F7435, MilliporeSigma) at 1:100 diluted superblock at 4°C for 1 h. The flag tag on these constructs is on the N-terminus, and thus can only be labeled in live cells if the OR reaches the cell surface. Cells were then washed, fixed with 4% paraformaldehyde (PFA), permeabilized with 0.3% Triton X-100, and blocked with diluted superblock for 1 h at room temperature. Total expression staining was assayed in fixed, permeabilized cells by staining with mouse monoclonal anti-flag antibody (M2, Cat. No. F1802, MilliporeSigma) at 1:100 in diluted superblock at 4°C overnight. All surface and total staining cells were incubated with secondary antibody at 1:1000 in diluted superblock with Hoechst 33342 (Cat. No. LSH3570, Invitrogen Molecular Probes) at 1:2500 dilution in the dark for 1 h at room temperature. Finally, cells were washed and mounted using VectaShield Hard Set mounting medium (H-1400, Vector Laboratories). Images were taken using a confocal microscope (Zeiss LSM 700). Images were quantified manually by ImageJ by quantifying the signal per cell (quantified in every positive cell within the field of view; values for cells within the same field of view were averaged). Thus, each “n” represents the average of values for a given field of view.

### Real-time cAMP assay

Human embryonic kidney 293T (HEK 293T) cells were seeded into a poly-L-lysine coated black 96-well-plate with a clear bottom and incubated overnight. Cells were transfected with OR51E1 or OR51E1 variants with RTP1S and/or Golf. After 4 h transfection, the cADDis cAMP sensor (BacMan baculovirus, Cat. No. U0205G, Montana Molecular) was transduced into the cells. The cAMP fluorescence was measured 24 h after the sensor transfection at excitation 490 nm and emission 525 nm wavelengths. Real-time cAMP production was measured before and every minute after stimulus for a total 10 min. The stimulus was presented during the 10 min sampling period. Stimuli were sodium butyrate (Cat. No. 303410, MilliporeSigma), isovaleric acid (IVA, Cat. No. 129542, MilliporeSigma), valeric acid (VA, Cat. NO. 240370, MilliporeSigma), 3-methylvaleric acid (3-MVA, Cat. No. 222453, MilliporeSigma), 4-methylvaleric acid (4-MVA, Cat. No. 277827, MilliporeSigma) and cyclobutane-carboxylic acid (CCA, Cat. No. C95609, MilliporeSigma).

### ELISA

Total protein expression of the OR51E1 constructs in HEK 293T cells was measured by ELISA, as previously reported^45,46^. Cells were grown in a poly-L-lysine 96-well-plate (black plate with clear bottoms) and transfected with OR51E1 or OR51E1 variants with RTP1S, and with or without Golf. After 24 h transfection, cells were fixed, permeabilized, and incubated with mouse monoclonal anti-flag antibody (M2, Cat. No. F1802, MilliporeSigma) at 1:100 dilution and detected with anti-mouse horseradish peroxidase (HRP)-conjugated secondary antibody (Cat. No. 115-035-146, Jackson) at 1:8000 dilution. HRP levels were measured with 1-Step Ultra TMB (3,3’,5,5’-tetramethylbenzidine; Cat. No. 34028, ThermoFisher Scientific) at excitation of 450 nm.

### Statistical analysis

Data are presented as mean ± standard error of the mean (SEM). Sample size (n) is indicated for each reported value. Statistical significance was determined by t-test, or, ANOVA for multiple groups (*p*<0.05). For human genetic data analysis, linear regression analyses were performed using R statistical software^47^.

## Results

### Olfr558 is expressed in vascular smooth muscle cells

Olfr558 is expressed in the olfactory epithelium (**Fig. S1**) as well as in non-nasal tissues. We previously reported that Olfr558 is expressed in the kidney^33^. RNAScope staining in the kidney revealed that Olfr558 is expressed in renal vascular smooth muscle cells (co-stained by an antibody for α-smooth muscle actin, α-SMA) in Olfr558 wild-type (Olfr558 WT) but not Olfr558 knockout (Olfr558 KO) kidneys (**Fig. 1A**). Olfr558 is also expressed in both afferent and efferent arterioles in the kidney, as identified by co-staining for α-SMA adjacent to a glomerulus (**Fig. 1B**) as well as co-staining for renin (**Fig. 1C**).

**Fig. 1.**
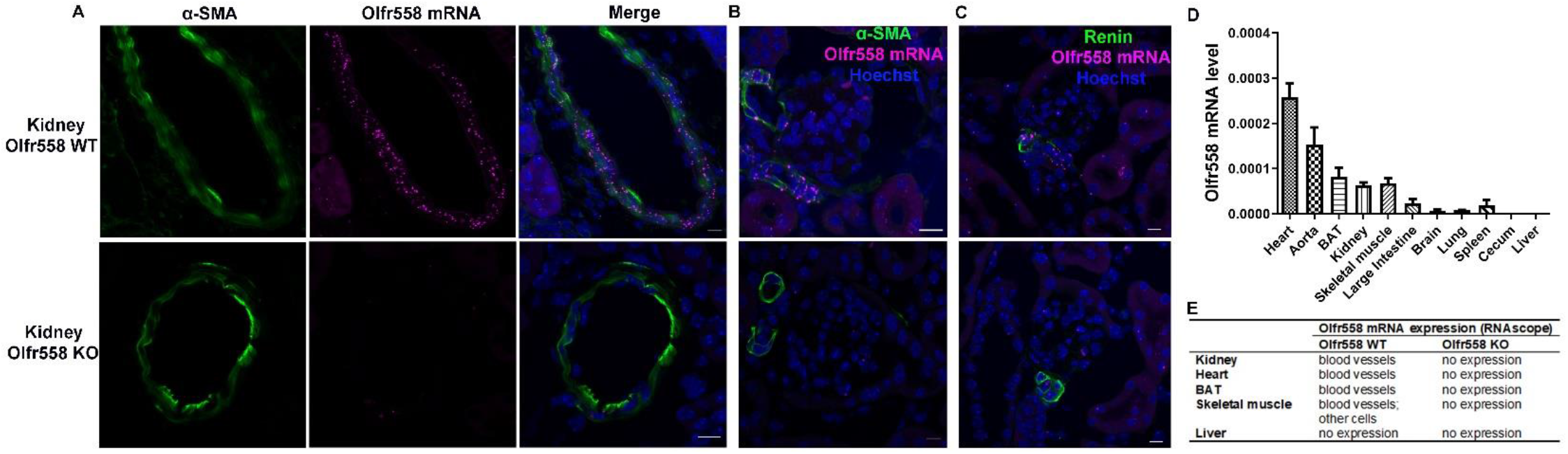
Olfr558 is expressed in vascular smooth muscle cells. **(A)** Immunostaining (α-SMA, green, blood vessel marker) and RNAScope (Olfr558 mRNA, purple) demonstrate that Olfr558 localizes to vascular smooth muscle cells in Olfr558 WT kidney (Hoechst nuclear stain, blue, in merge). Olfr558 mRNA staining is absent in Olfr558 KO kidney. **(B)** Olfr558 mRNA is expressed in both afferent and efferent arterioles in Olfr558 WT, but not Olfr558 KO, kidney (Hoechst nuclear stain, blue). **(C)** Co-staining with a renin antibody (green) and Olfr558 RNAScope probe (purple) demonstrates that Olfr558 is expressed in renin positive cells in Olfr558 WT, but not Olfr558 KO, kidney. Scale bar: 10 μm. **(D)** Olfr558 expression in a subset of tissues by qPCR. Data normalized to GAPDH and averaged from n=3 males and 3 females. **(E)** Summary of Olfr558 localization in kidney, heart, BAT, skeletal muscle, and liver.

To determine if the localization of Olfr558 to blood vessels occurs beyond the kidney, qPCR was used to identify other organs which express Olfr558 (**Fig. 1D**), and subsequently RNAScope was performed on tissues with relatively high expression (**Fig. 1E; Fig. S2-4**). By qPCR, Olfr558 is relatively highly expressed in the heart, aorta, brown adipose tissue (BAT), kidney, and skeletal muscle, but exhibits little to no expression in the large intestine, brain, lung, spleen, cecum or liver. In heart, BAT, and skeletal muscle, Olfr558 localized primarily to vascular smooth muscle by RNAScope, although there is rare expression in a minority cell type in skeletal muscle. We noted that Olfr558 is typically expressed in some, but not all, vascular smooth muscle cells in a given vessel cross-section. The closest ‘sibling’ receptor to Olfr558, Olfr78, is also expressed in a subset of vascular cells in renal blood vessels^48^. In olfactory neurons, the expression of olfactory receptors is thought to be largely mutually exclusive (one cell-one receptor rule)^49^. To determine if Olfr78 and Olfr558 are co-expressed in the kidney, we performed RNAScope for both receptors. We find that the majority of cells express only Olfr558 (47%) or Olfr78 (37%), but 16% of cells co-express both receptors (**Fig. S5)**.

### Olfr558 is required for sex differences in blood pressure

Olfr558 WT and KO mice exhibit no genotypic differences in body weight (**Fig. S6A**), kidney weight/body weight ratio (**Fig. S6B**), heart weight/body weight ratio (**Fig. S6C**), or glomerular filtration rate (GFR) (**Fig. S6D**). Likewise, there are no genotypic differences in blood electrolytes, glucose, blood urea nitrogen (BUN), creatinine, or hemoglobin (**Supplementary Table 1**). Based on the localization of Olfr558 to vascular smooth muscle cells, we hypothesized that Olfr558 may play a role in blood pressure regulation. To test this hypothesis, we used telemetry to measure 24-hour blood pressure. It is well-established that males have higher blood pressure than premenopausal females by ∼10mmHg in both humans^2-14^ and mice^1^. In agreement with this, we find that Olfr558 WT males exhibit significantly higher blood pressure than WT females, including mean arterial pressure (MAP) (**Fig. 2A**), systolic blood pressure (SBP) (**Fig. 2B**) and DBP (**Fig. 2C**). However, sex differences in blood pressure are absent in Olfr558 KO (**Fig. 2D-F**). Olfr558 KO females have a significant increase in MAP and DBP in both light (**Fig. 2 G, I**) and dark cycles (**Fig. 2 J, L**) as compared to WT females. Olfr558 KO males, as compared to WT males, exhibit a decrease in DBP without a change in SBP; the drop in DBP is statistically significant during the light cycle (**Fig. 2 G-I**). There are no genotypic differences in heart rate, although female KOs do have higher heart rates than male KOs during the light cycle (**Fig. S7A-C**). No genotypic or sex differences were seen in pulse pressure (**Fig. S7E-H**). Of note, other sex differences (not related to blood pressure) are intact in Olfr558 KO: for example, KO males weigh more than KO females (**Fig. S6A**), and KO males and females are fertile. Given the lack of sex differences in blood pressure in KO mice, we expanded our localization of Olfr558 to include testes and ovaries: in testes, Olfr558 is expressed primarily in blood vessels, but is also seen in a minority cell type (**Fig. S8**). In ovaries, Olfr558 is solely expressed in blood vessels (**Fig. S9**).

**Fig. 2.**
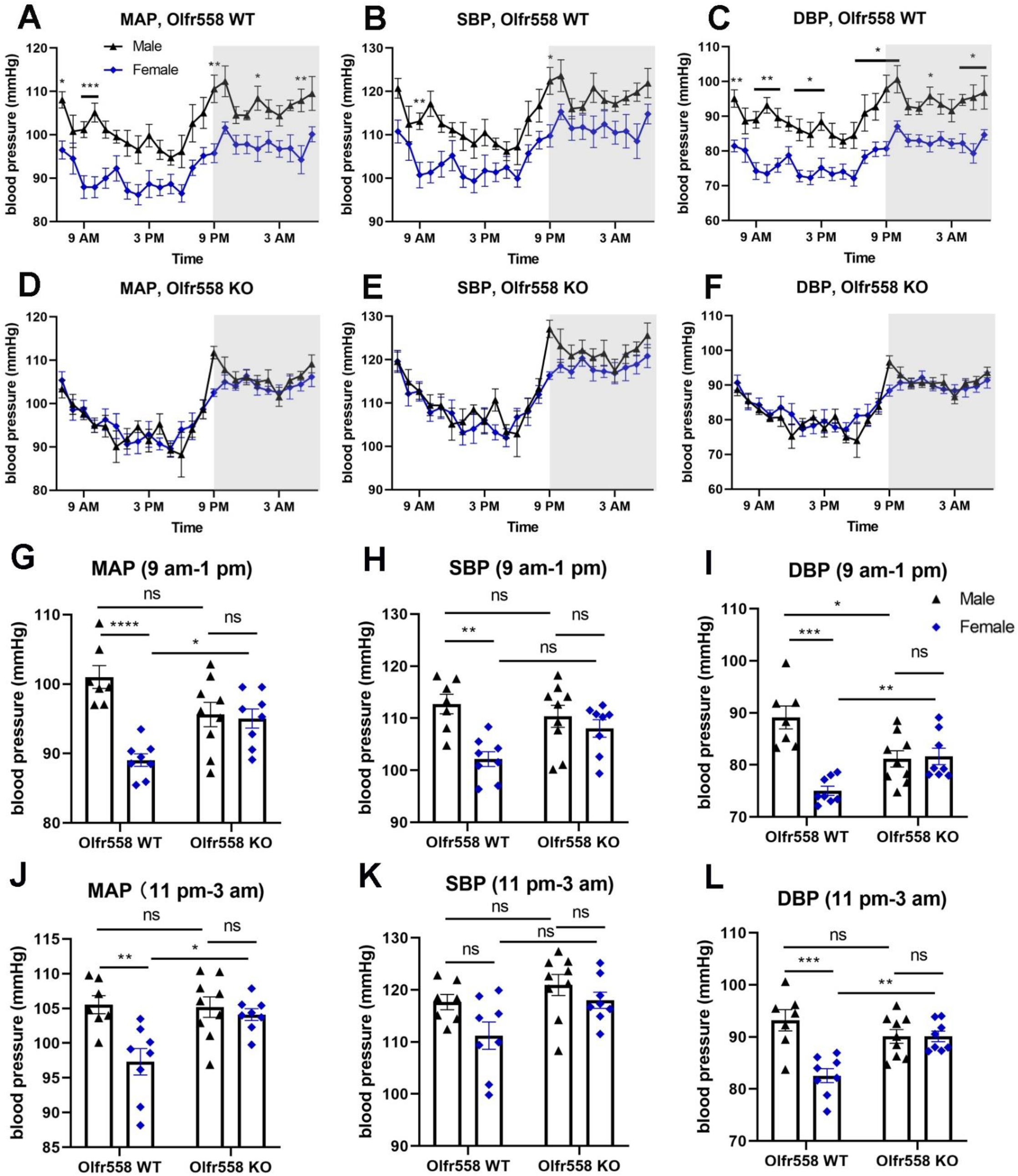
Olfr558 regulation of blood pressure is sex-specific. Blood pressure was measured by telemetry. Olfr558 WT mice exhibit previously reported sex differences in **(A)** MAP, **(B)** SBP, and **(C)** DBP. These differences are absent in Olfr558 KO **(D-F)**, due to increased blood pressure in females, along with decreased male DBP. Grey areas depict the active (dark, 9pm-7am) phase, and white areas depict the resting (light, 7am-9pm) phase. Averaged data from the light cycle (9am-1pm) are shown in **(G-I)**, and averaged data from the dark cycle (11pm-3am) are shown in **(J-L)**. Male WT: n=7, male KO: n=9. Female WT and KO: n=8. \**p*<0.05, \*\**p*<0.01; \*\*\**p*<0.001, \*\*\*\**p*<0.0001 by phenotype (Olfr558 WT vs. KO) and sex (Male vs. Female) using two-way ANOVA. ns: non-significant.

### Olfr558 modulates renin in males

Given the localization of Olfr558 to renin-expressing cells in the renal afferent arteriole, we examined a potential role for Olfr558 to regulate renin. Using qPCR, we find that Olfr558 mRNA is absent in KO kidneys (**Fig. 3A, 3F**), as expected. In female KO kidneys, we find no differences in renin mRNA (**Fig.3B**), no differences in the percentage of renin positive to total glomeruli (**Fig. 3C, D**), and no differences in plasma renin activity (PRA) (**Fig. 3E**). In male KO kidneys, we find a significant decrease in renin mRNA (**Fig. 3G**), a reduction in the percentage of renin positive to total glomeruli (**Fig. 3H**), and decreased PRA on two different diets (0.2%Na, **Fig. 3I**; 0.49% NaCl, **Fig. 3J**). These data suggest that Olfr558 acts to support renin levels in WT males, and thus changes in renin may contribute to the lower blood pressure in male KO.

**Fig. 3.**
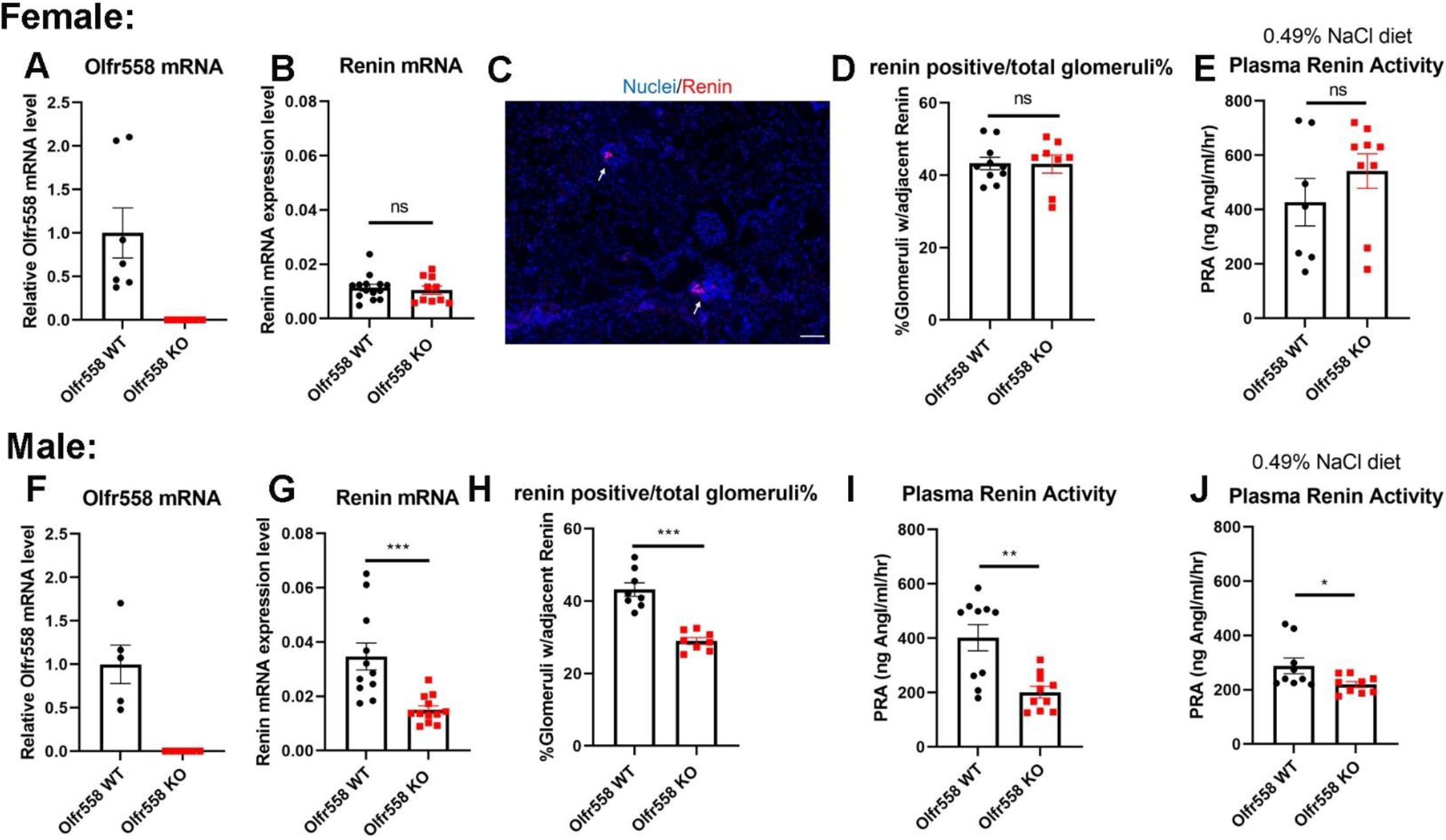
Olfr558 modulates renin in males. **(A, F)** Olfr558 mRNA is absent in kidneys of Olfr558 KO. Renin mRNA levels in the kidney are not altered in females **(B)**, but are decreased in KO males **(G)**. Renin staining (red, white arrow) is observed in glomerulus **(C)**, scale bar: 50 μm. **(D, H)** The ratio of renin positive/total glomeruli is reduced in male KOs, but not females. Plasma renin activity (PRA) is decreased in male KOs on standard chow **(I)** and 0.49% NaCl diet **(J)**, but is not altered in KO females **(E)**. The relative mRNA expression is presented normalized to its GAPDH or GAPDH and WT of the same sex. \**p*<0.05, \*\**p*<0.01, \*\*\**p*<0.001, ns: non-significant by t-test.

To determine if the changes in male KO are due to a decreased number of renin cells, we co-stained renin and α-SMA in male Olfr558 WT and KO kidneys (**Fig. 4A**), and quantified the number of cells that co-stained for both markers, or, for either one alone (as done previously^50^). We find a decreased number of cells staining for ‘renin only’ in male KO mice, and to a lesser extent, a decreased number of cells labeling for both α-SMA and renin (**Fig. 4B**). Similarly, qPCR for the renin cell marker Akr1b7 revealed a decrease in renal Akr1b7 expression in male but not female KO (**Fig. 4C, D**).

**Fig. 4.**
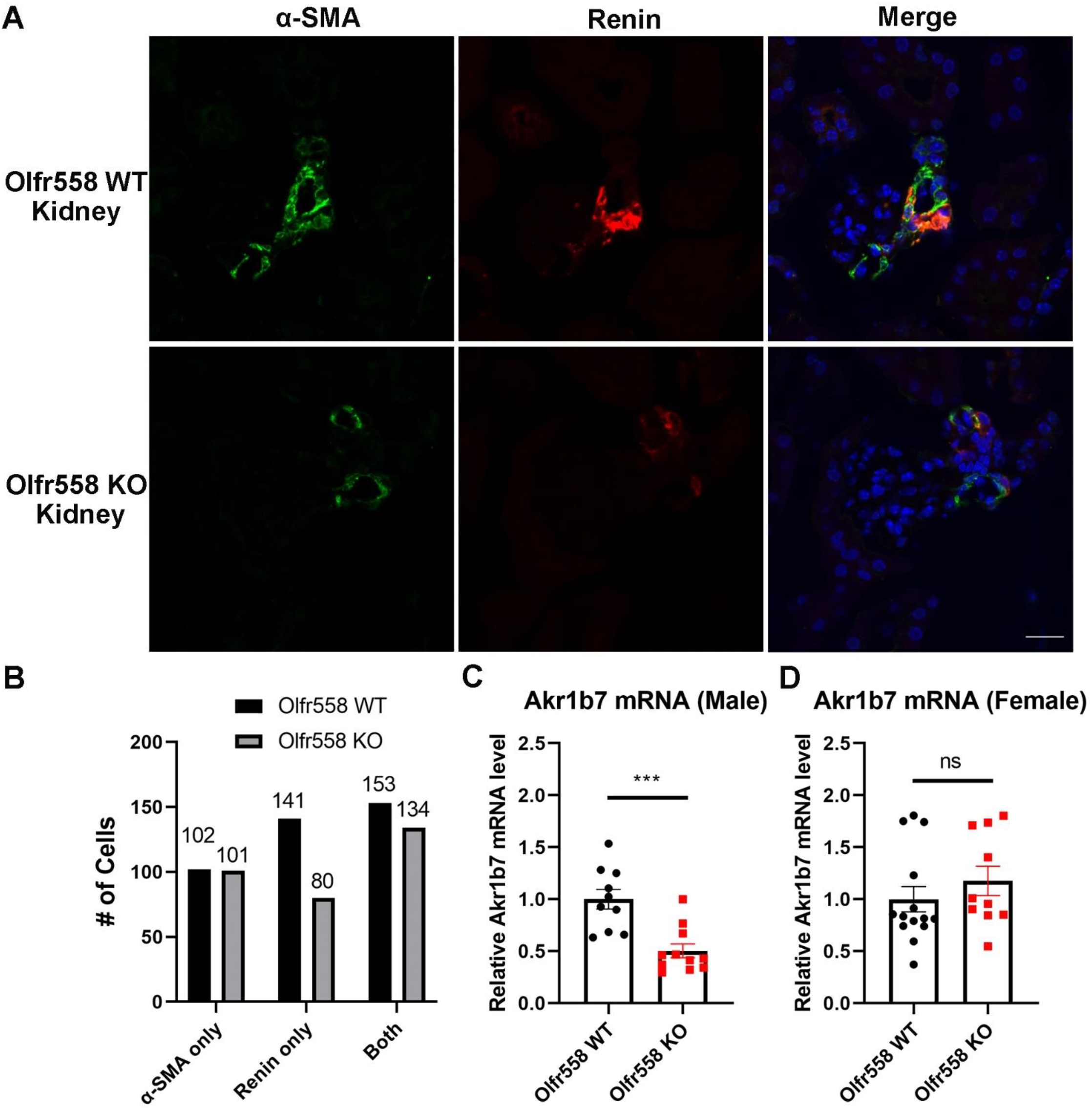
Renin expressing cells are decreased in male Olfr558 KOs. Co-staining of α-SMA (green) and renin (red) was examined from male WT and KO kidneys **(A)**. JGA-associated cells were counted as α-SMA only, renin only, or α-SMA and renin (both) expression; this quantification indicates that the number of renin-expressing cells is decreased in male KO **(B)**. n=4 WT and KO. Akr1b7 (renin cell marker) mRNA expression is reduced in KO males **(C)** but not females **(D)**. Scale bar: 20 μm. \*\*\**p*<0.001, ns: non-significant by t-test.

### Olfr558 modulates vasoreactivity in males

Given the localization of Olfr558 to vascular smooth muscle cells in a variety of vascular beds, we examined a role for Olfr558 to regulate vasoreactivity in aortic rings and mesenteric arteries. *Ex vivo* vascular reactivity studies show that Olfr558 KO aortic rings from females exhibit higher maximal contractility induced by potassium chloride (KCl) as compared to WT. Similar responses to phenylephrine (PE), acetylcholine (ACh), and sodium nitroprusside (SNP)(**Fig. 5A-D**), suggest no impairments in vascular smooth muscle or endothelial function in female KO. In males, KO aortic rings exhibit less constriction to PE (**Fig. 5F**), which could contribute to the hypotension seen in these mice. In contrast, there were no genotypic differences in males to the constriction to KCl (**Fig. 5E**) or relaxation to ACh (**Fig. 5G**) or SNP (**Fig. 5H**).

**Fig. 5.**
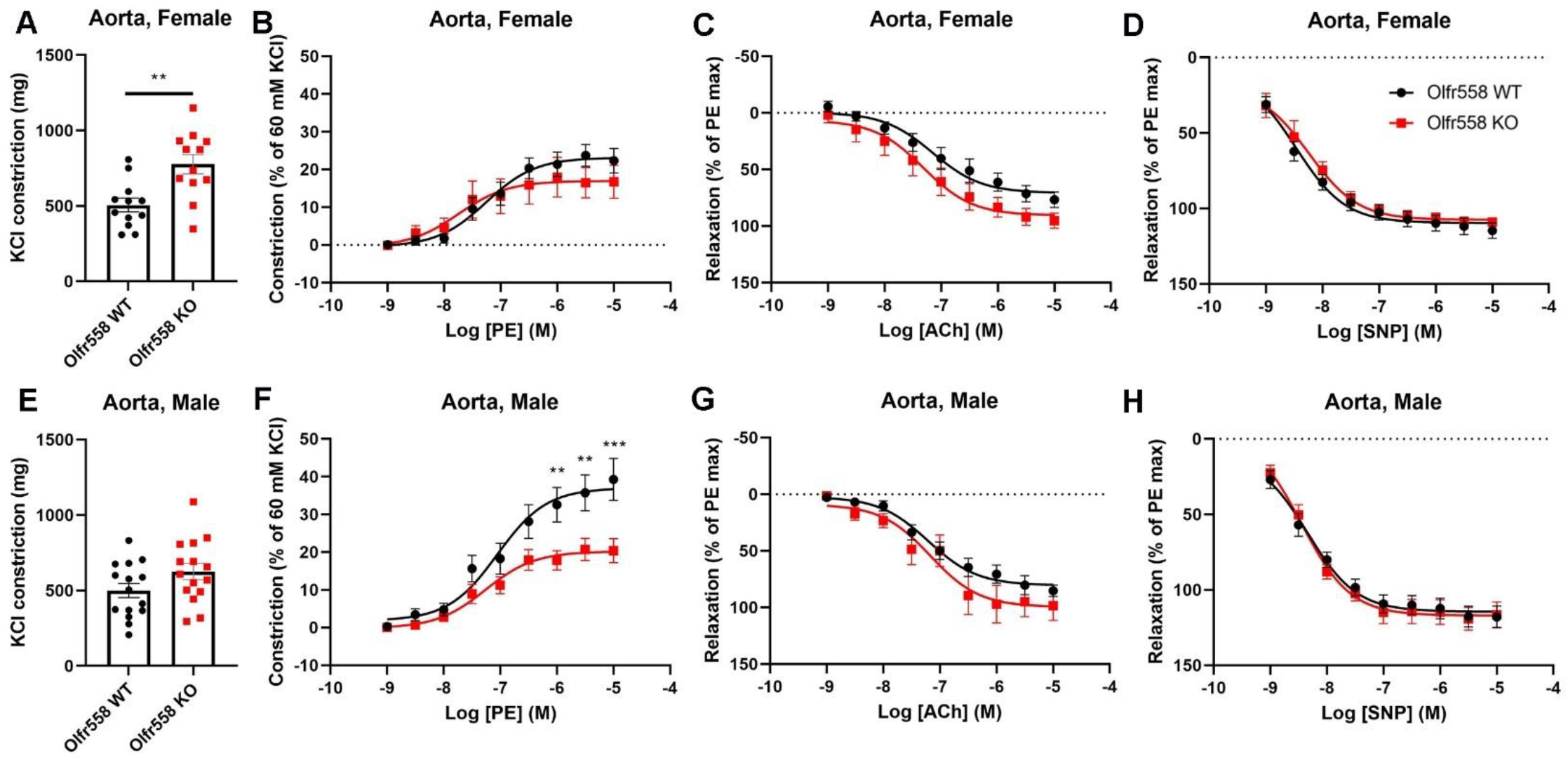
Aortic vasoconstriction to PE is impaired in Olfr558 KO males. Aortic constriction to 60 mM KCl was enhanced in females **(A)** but not males **(E)**. Aortic constriction to increasing doses of PE was enhanced in males but not females **(B, F)**. There were no genotypic changes in relaxation to ACh (endothelium-dependent, **(C, G)**) or to SNP (endothelium independent, **(D, H)**). PE responses are normalized to the KCl response, and ACh and SNP responses are normalized to the maximum PE response. n=12∼15 aortic rings from n=6 Olfr558 WT and KO males. \*\**p*<0.01, \*\*\**p*<0.001 by t-test for A and E, two-way ANOVA for B-D and F-H.

Olfr558 KO mesenteric arteries from females demonstrate similar constriction to KCl and PE, and similar relaxation to ACh and SNP (**Fig. 6A-D**). In males, there are no genotypic differences in the constriction to KCl and PE, or in the relaxation to ACh (**Fig. 6E-G**). However, Olfr558 KO mesentery from males exhibit increased relaxation to SNP, which could contribute to the decreased blood pressure in males (**Fig. 6H**).

**Fig. 6.**
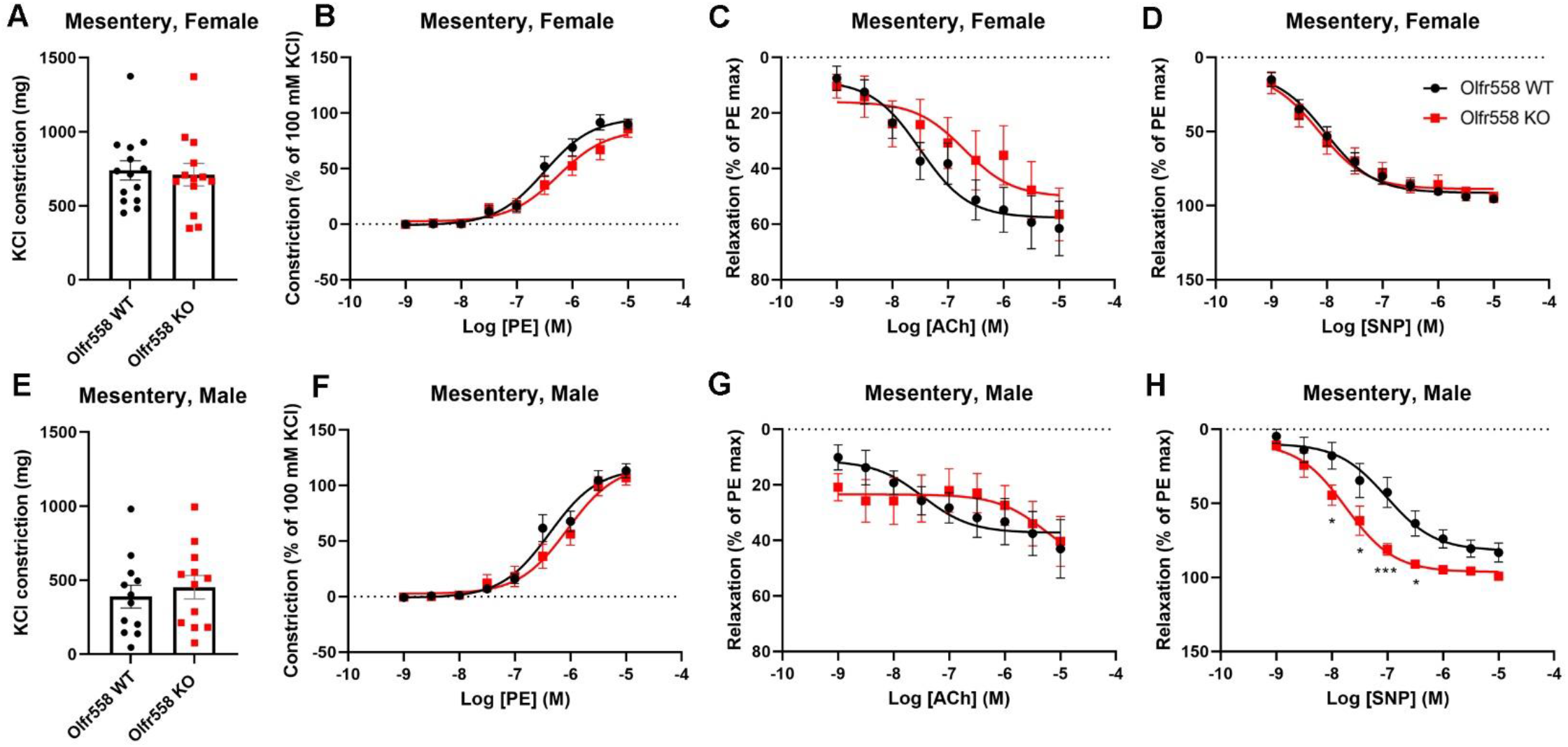
Mesentery vasorelaxation to SNP is enhanced in Olfr558 KO males. No genotypic differences were observed in mesentery constriction to 100 mM KCl **(A, E)**, mesentery constriction to PE **(B, F)**, or mesentery relaxation to ACh **(C, G)**. However, endothelium-independent vasorelaxation in response to SNP was enhanced in KO males (**H**) but not females **(D)**. PE responses are normalized to the KCl response, and ACh and SNP responses are normalized to the maximum PE response. n=8∼14 Mesenteric arteries from n=7 Olfr558 WT and KO females. \*\**p*<0.01, \*\*\**p*<0.001 by t-test for A and E, two-way ANOVA for B-D and F-H.

### Olfr558 modulates arterial stiffness in females

Data from the studies described above imply that the decreased DBP in the males may be driven by decreased renin, lessened constriction to PE in the aorta and/or exaggerated relaxation to SNP in the mesentery. However, the only genotypic difference in females was KCl constriction in the aorta. Thus, to better understand the female phenotype, we evaluated the elastic properties of female aortas *ex vivo* and measured arterial stiffness in females *in vivo*. The tensile properties are similar between Olfr558 WT and Olfr558 KO in both intact and decellularized aortic rings from 10-week-old female mice (**Fig. 7A**). Likewise, there are no differences in the wall thickness and diameter of aortic rings from 10-week-old Olfr558 WT and KO females (**Fig. 7B and C**). However, pulse wave velocity (PWV) is significantly increased in 10-week-old Olfr558 KO females as compared to WT (**Fig. 7D**), implying that increased vascular stiffness contributes to the female KO phenotype. Taken together, these data suggest that the net *in vivo* smooth muscle tone is responsible for the higher PWV noted in KO females, but that Olfr558 does not contribute significantly to passive mechanical stiffness of the matrix.

**Fig. 7.**
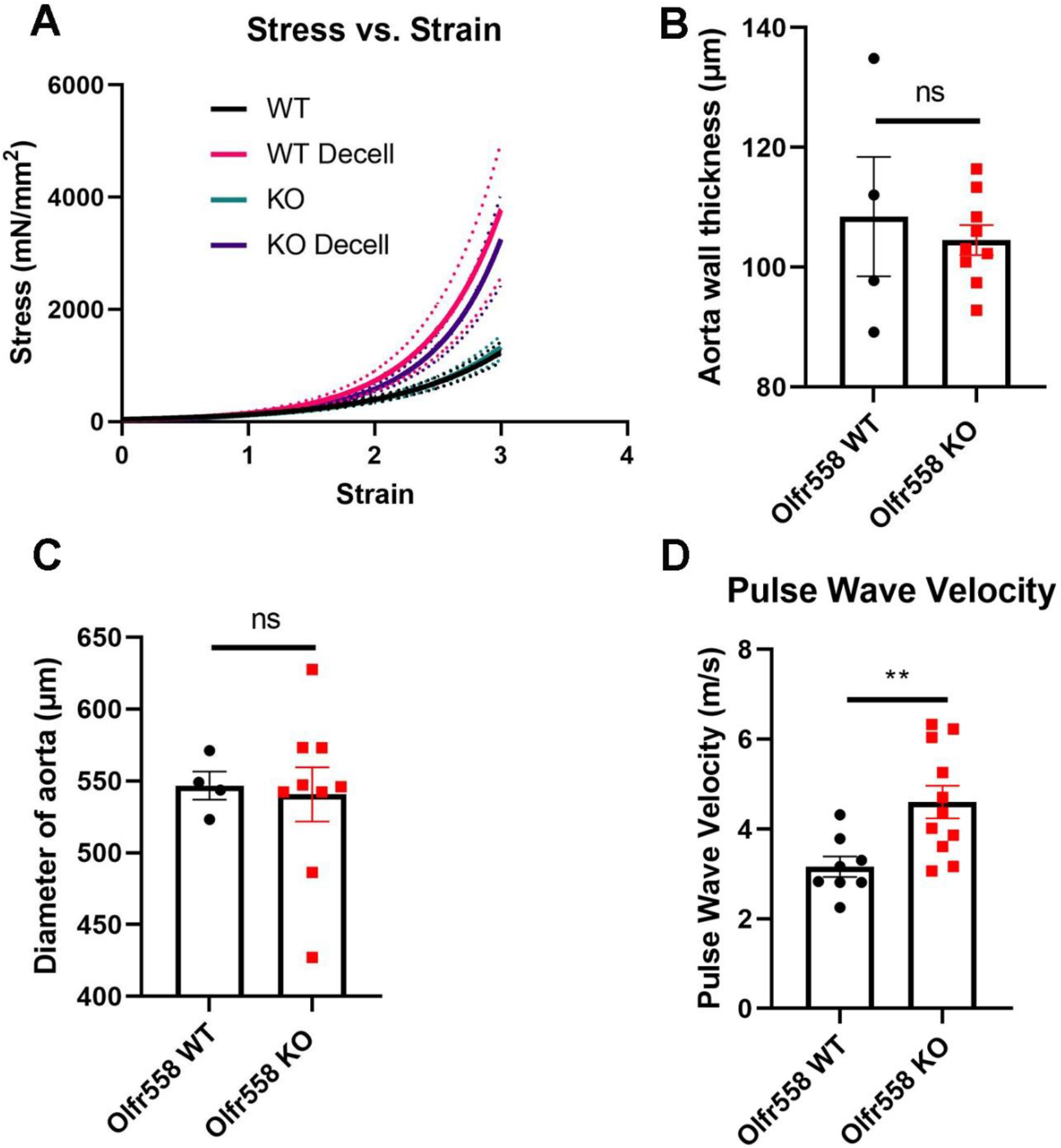
Increased arterial stiffness in Olfr558 KO females. **(A)** Tensile testing of intact and decellularized thoracic aortic rings from Olfr558 WT and KO females did not reveal genotypic differences. Likewise, there were no genotypic differences in wall thickness **(B)** or aortic diameter **(C)**. (A-C: n=4 for WT, n=9 for KO) **(D)** However, pulse wave velocity, an *in vivo* measure of arterial stiffness, is higher in Olfr558 KO females. n=8 for WT, n=11 for KO. **p<0.01 by t-test, ns: non-significant.

### Gut microbiome is similar in Olfr558 WT and KO mice

Butyrate, a short chain fatty acid (SCFA) produced by gut microbes, is the ligand which yields the strongest activation of both Olfr558 and OR51E1^46,51^.To determine if alternations in the gut microbiome of Olfr558 KO mice may contribute to the phenotype, we analyzed 16S rRNA from fecal DNA from Olfr558 WT and KO (females and males). The major phyla show no differences between WT and KO (**Fig. S10**).

### Human Genetic Analysis

The human ortholog of Olfr558 is OR51E1. An OR51E1 variant was previously identified by Evangelou, et al^44^ as significantly associated with diastolic blood pressure (rs17224476): in the total GWAS meta-analysis (N∼750k), each copy of the minor allele A increased DBP by ∼0.16 mmHg (*p*=7.37× 10^−9^) according to an additive allelic association model. However, this previous GWAS did not examine sex-specific effects. So, here, we performed sex- and age-stratified analyses on OR51E1 variants using UKB data. Our analysis focused on the lead, common, missense variant (rs17224476) from the Evaneglou, et al study, and, also on a rare missense variant (rs202113356) which we have previously studied in vitro^46^. Within UKB data, the common variant (rs17224476) has Minor Allele Frequency (MAF) = 11.1%, and the rare variant (rs202113356) has MAF = 0.025%. Even though these two variants are located very close together on the genome, only 434 bp apart, they are independent, uncorrelated variants, and calculated within UKB to not be in Linkage-Disequilibrium (r^2^ ∼1 × 10^−5^, very close to 0).

We confirm the previously published association of the lead variant in the total sample, with the association for DBP still being highly significant in the UKB data alone (*p*=4.27 × 10^−5^; N = 423,655). This lead variant is significantly associated with DBP within each of the separate men and women subgroups with the minor allele A increasing DBP similarly in both men and women, showing no significant sex-interaction (*p*=0.76) (**Supplementary table 2, Fig. 8A-C**). This variant is also significant (*p*<0.05) for females<50, females>50 and males >50 within the stratified analyses (**Fig. 8A**), and with no significant evidence of any age-interaction effect (**Fig. 8C**). So, for this common variant there is sufficient statistical power to clearly demonstrate the significant association of *OR51E1* with DBP overall, showing across all subgroup analyses, that the minor allele A of this variant increases DBP. These data therefore confirm and expand the role of this OR in blood pressure regulation, and imply that common frequency variants in this receptor affect blood pressure equally in men and women.

**Fig. 8.**
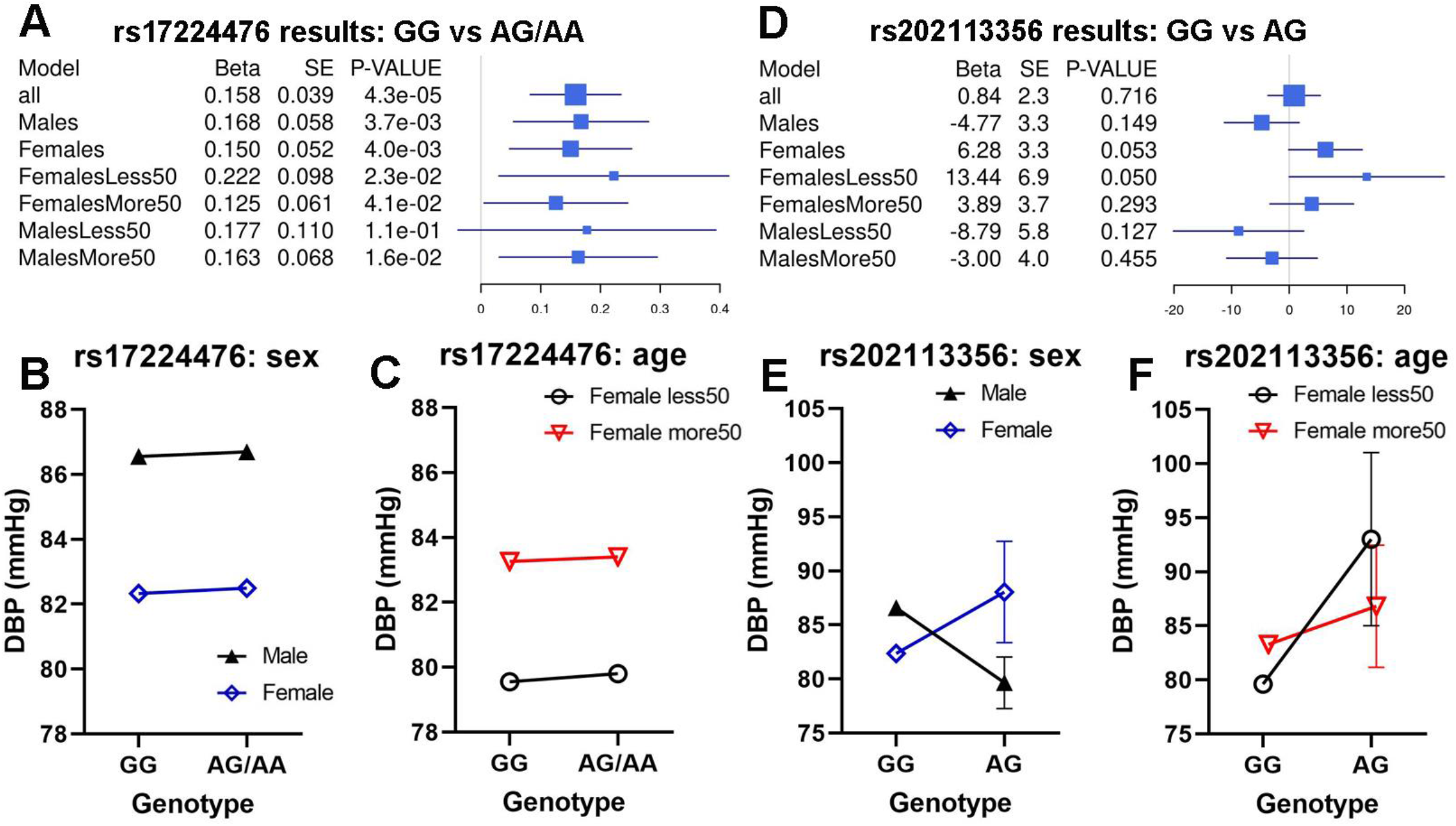
OR51E1 variants and blood pressure. **Figures A-C** are or the lead variant (rs17224476); **Figures D-F** are for the rare variant (rs202113356). **Figures A** & **D** show forest plots illustrating all association results for different subgroup models for each variant. A positive beta value for the linear regression effect estimate indicates that the minor allele A of the variant increases DBP, whereas a negative beta value indicates that the minor allele A of the variant decreases DBP. *p*<0.05 is declared as significant. **Figures B** & **E** illustrate the sex-stratified analyses, comparing the mean DBP in non-carriers (GG) vs carriers of the minor allele (AG/AA) separately in males vs females. **Figures C** & **F** illustrate the age-stratified analyses, comparing the mean DBP in non-carriers (GG) vs carriers of the minor allele (AG/AA) separately in younger women <50 years vs older women > 50 years. In these stratified analysis plots, the mean DBP values within each subgroup are plotted together with the interval limits for the Standard Error of the Mean (SEM) bars. The lead OR51E1 variant (rs17224476) is significantly associated with DBP overall and within sex- and age-stratified analyses **(A-C)**. The minor allele A of this variant increases DBP within all individuals, without any sex-**(B)** or age-**(C)** interactions. The rare OR51E1 variant (rs202113356) has a significant sex-interaction effect (*p*=0.0148) **(E)**, with sex-specific, opposing effects on DBP in males vs females **(D-F)**. The rare variant decreases DBP in men but increases DBP in women **(D)**

We also examined a rare variant which is also localized to an exon and is primarily seen in South Asia. We previously published that this rare variant alters OR51E1 function *in vitro* (rs202113356)^34^. The highlighting result from our UKB analyses is that this rare variant has a significant sex-interaction result (*p*=0.015), showing sex-specific effects, with opposite directions of effect in men vs women (**Supplementary table 2, Fig. 8D-F**). The association is nominally significant (*p*=0.053) within females, with the minor allele A increasing DBP by +6.28mmHg for female carriers (AG) vs female non-carriers (GG). In contrast, within males, the effect estimate (−4.77mmHg) indicates that the rare variant decreases DBP in men, and the association result is not significant (*p*=0.149), which could either be due to a lack of association in males completely relating to the sex-specific nature of the variant, or simply due to lack of statistical power for analysis of the rare variant. (**Fig. 8D**). Furthermore, from the age-stratified analyses within females, there is a stronger association in younger females with a more significant result, despite a lower sample size (*p*=0.0501; N=57,464) in women <50 years compared to in women >50 years (*p*=0.29; N=171,067), and also a much larger effect size (+13.44mmHg in women <50 vs +3.89mmHg in women >50) (**Fig. 8D**), as can also be seen by the steeper gradient of the effect slope (**Fig. 8F**), although there is no significant age-interaction effect (*p*=0.26), likely due to a lack of statistical power for detecting a higher-order interaction effect within a smaller stratified analysis sample for a rare variant. Acknowledging that we always expect less statistical power for such a rare variant, the significant sex interaction results for this rare variant (**Fig. 8E**) is therefore a very clear conclusion, indicating that this variant has a strong, differential effect on DBP with opposing effects in men versus women.

### OR activity is reduced in OR51E1 variants transfected cells

Given these findings, we further examined both variants using *in vitro* studies. The common (rs17224476) variant results in a missense change, (S10N). (Of note, the “S” residue is conserved in numerous species, including humans, mouse, and rat.) We find that OR51E1 S10N shows similar cell surface trafficking (**Fig. S11**) and total protein expression (**Fig. S12**) as wild-type OR51E1. However, in the absence of Golf, butyrate induced cAMP production is significantly and dose-dependently attenuated in OR51E1 S10N transfected cells (**Fig. 9A and B**). Results with other ligands (which we have previously identified for the wild-type receptor^33^) did not reach statistical significance in the absence of Golf (**Fig. 9C**). When co-transfected with Golf, the reduction in OR51E1 S10N signaling is significant for all ligands tested (**Fig. 9B, D**).

**Fig. 9.**
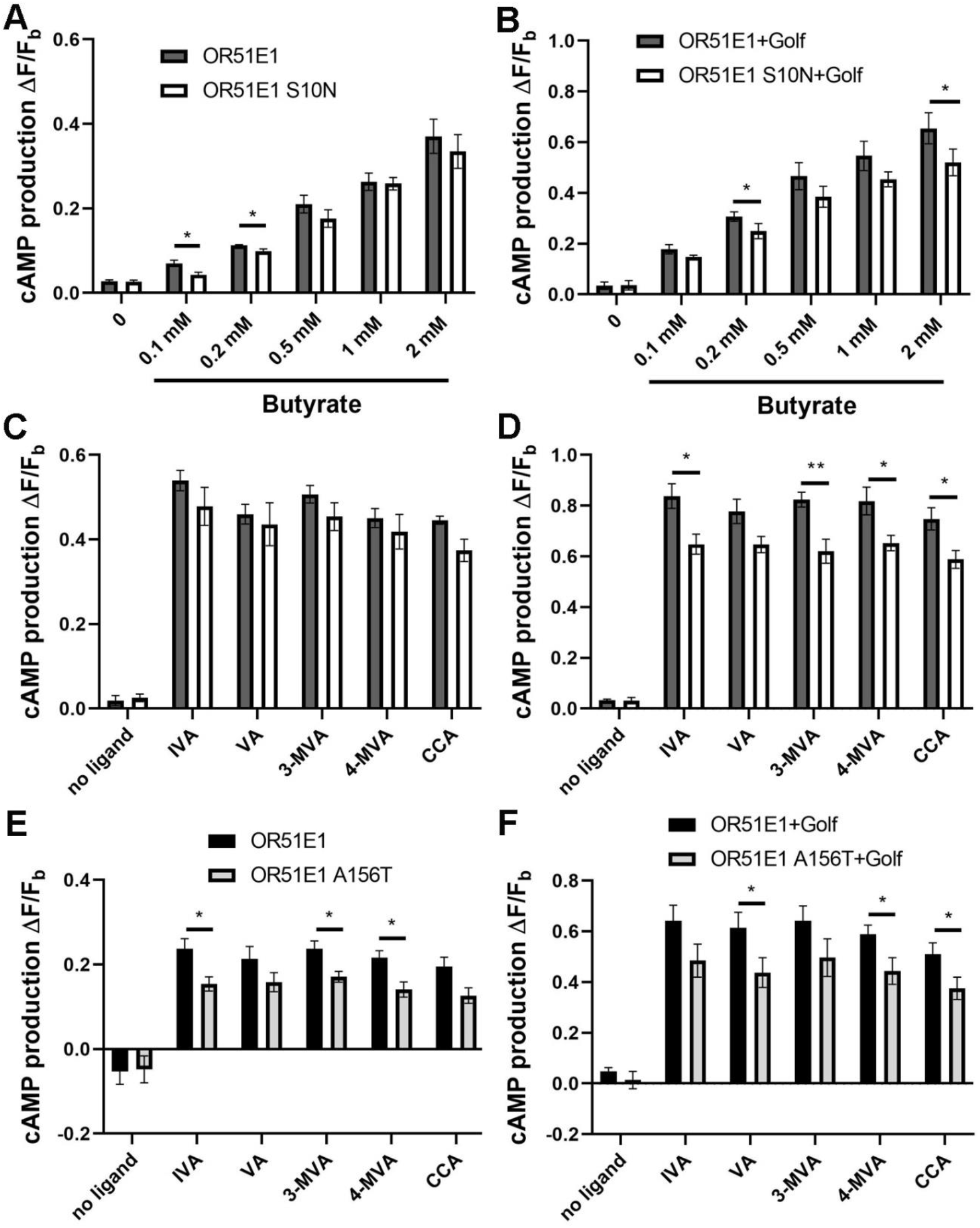
OR51E1 S10N has reduced activity as compared to wild-type OR51E1. (**A-B**) Butyrate-induced cAMP production is significantly attenuated in OR51E1 S10N transfected cells with or without Golf co-transfection. For other ligands, the difference between wild-type and S10N is significant with but not without Golf (**C-D**) (**D**). (**E-F**) Ligand-induced cAMP production is reduced in OR51E1 A156T transfected cells with or without Golf co-transfection. IVA: isovaleric acid, VA: valeric acid, 3-MVA: 3-methylvaleric acid, 4-MVA: 4-methylvaleric acid, CCA: cyclobutane-carboxylic acid. Ligands in C-D were tested at 1mM. \**p*<0.05, \*\**p*<0.01 OR51E1 (+Golf) vs. OR51E1 S10N or OR51E1 A56T (+Golf) by ANOVA.

The rare variant (rs202113356) also results in a missense change (A156T). For OR51E1 A156T, we recently published that OR51E1 A156T exhibits less cell surface but similar total protein expression as OR51E1^46^. We also reported that butyrate activation is significantly and dose-dependently reduced in OR51E1 A156T and/or Golf transfected cells^46^. Here, we further evaluated the response of OR51E1 A156T to other ligands, and found that cAMP production is also attenuated in OR51E1 A156T transfected cells for all ligands tested (**Fig. 9E, F**).

## Discussion

We report here that Olfr558 is required for sex differences in blood pressure. Olfr558 localizes to vascular smooth muscle cells in a subset of non-nasal tissues including kidney, heart, BAT, skeletal muscle, testes, and ovaries, but is absent from other tissues (e.g., liver). In the kidney, Olfr558 is expressed in both afferent and efferent arterioles, including renin positive cells. Functional studies using Olfr558 WT and KO mice demonstrate that Olfr558 is required for sex differences in blood pressure. Olfr558 KO males exhibit decreased DBP, whereas Olfr558 KO females have increased MAP and DBP. Lowered DBP in the males may be driven by decreased plasma renin, a muted constriction to PE in the aorta, and an exaggerated relaxation to SNP in the mesentery. The increased blood pressure in KO females is likely driven by increased vascular stiffness, as indicated by the elevated pulse wave velocity. In humans, OR51E1 has been identified as a locus that is associated with significant changes in DBP. We find that a rare OR51E1 variant induces changes in DBP which match both the direction (increased in females, and decreased in males) and the magnitude seen in KO mice of each sex. Together, these data imply that this evolutionarily conserved OR is required for sex differences in blood pressure regulation through a mechanism that involves modulation of both renin and *in vivo* smooth muscle tone.

### Olfr558 and OR51E1 Localization

Our studies demonstrate that Olfr558 is broadly expressed in non-nasal tissues, where it localizes to blood vessels. Our findings are supported by a report that Olfr558 is expressed in the blood vessels of the eye^24^, a report that the rat ortholog of Olfr558 (Olr63) is associated with glomeruli^52^, and a study which used microarray to identify genes that confer the identity of the renin cell^53^. Intriguingly, this latter study also reported that Olfr558 is upregulated in response to captopril treatment, implying that Olfr558 expression is regulated in response to changes in blood pressure. Finally, an RNAseq study examining human ectopic olfactory receptor expression found that OR51E1 is expressed in numerous tissues such as heart, kidney, adipose, ovary, testis, and skeletal muscle^31^, consistent with the expression profile we see in mice. Notably, we have not observed any differences in expression in males vs females. Finally, although there is a previous report that OR51E1 is expressed in renal epithelial cells (both in the HK-2 human proximal tubule cell line, and in human kidney)^54^, we did not observe localization to renal tubular epithelial cells by RNAScope.

### Sex differences in Blood Pressure

Sex differences in blood pressure in normotensive subjects are well-documented in both animal models^1^ and humans^1-14,55^. In humans, 24 hr ambulatory blood pressure measurements show that men have higher systolic blood pressure (∼10 mmHg) than women from the ages of 20-70; this difference is absent in those above the age of 70^55^. Furthermore, a study of 27,542 participants demonstrated that not only do women have lower blood pressure than men, but women incur increased risk for cardiovascular disease at lower blood pressure values compared to men^10^: for example, the stoke risk for women with SBP from 120-129mmHg is comparable to the risk for men with SBP from 140-149^10^, highlighting the clinical relevance of these sex differences. The sex differences in blood pressure seen in humans are also observed in mice, where the mean arterial pressure of males is ∼10 mmHg higher than that of females by telemetry^56^. This differences persists through 12 months of age but is absent by 18 months of age due to an increase in female blood pressure^56^. Consistent with previous report, we find that Olfr558 WT males have blood pressure which is ∼10 mmHg higher than females (Fig. 2). However, this blood pressure difference is absent in Olfr558 KO mice (Fig. 2), indicating that Olfr558 is required for sex differences in blood pressure. From our UKB analyses (Fig 8E), even though baseline blood pressure is higher in males, the sex-specific opposing effects of the rare variant (rs202113356) lead to male minor allele carriers having lower mean DBP than non-carrier females; and vice versa, that female minor allele carriers have higher mean DBP than non-carrier males, demonstrating how the effect of the genetic variant completely counterbalances and switches the normal underlying blood pressure levels in men and women.

### Renin

Sex differences in renin expression have previously been reported. For example, the renal renin concentration in adult females is lower than that of males in both Wistar and Sprague-Dawley rats^57^. Similarly, we find that Olfr558 WT female kidneys have less renin mRNA than male kidneys (Fig. 3B vs G, p<0.01 by t-test); this sex difference muted in KO mice. In our hands, plasma renin activity exhibited no differences between WT males and females (Fig. 3); this is in agreement with a previous study showing that PRA is similar in normotensive men and women^58^ as well as in Lewis rats^59^.

As for genotypic differences, we find that male KO (vs WT males) have decreased renin mRNA, decreased glomeruli-associated renin positive cells, and decreased PRA (Fig. 3 and 4). Of note, low renin paired with hypotension implies that the low renin is a contributing factor to the blood pressure change, as otherwise one would expect renin to be elevated with hypotension. However, if lowered renin were fully responsible for the male blood pressure phenotype diastolic and systolic pressure should be altered equally. Instead, KO males exhibit isolated diastolic hypotension. Furthermore, renin is not altered in females (Fig. 3). Therefore, Olfr558-induced sex differences in blood pressure cannot be explained by renin alone.

### Vascular Tone

We find that Olfr558 WT males and females exhibit similar responses to PE and ACh in both aortic rings and mesenteric arteries *ex vivo* (Fig. 5 and 6). This is consistent with a previous report that mesenteric arteries demonstrate similar relaxation to ACh in female and male rats, and were unaffected by ovariectomy^60^, but contradicts reports that aortic rings show more constriction to phenylephrine in male versus female rats^61,62^. It has also been reported that vascular constriction is significantly enhanced in ovariectomized versus intact female rats (with no difference between castrated and intact male rats), suggesting that the sex differences in vascular tone are likely related to sex hormones^63^. Of note, the maximum contraction to KCl in WT mesenteric arteries is higher in females vs males (Fig. 6).

As for genotypic differences, male KOs exhibit decreased constriction to PE in the aorta, and exaggerated relaxation to SNP in the mesentery as compared to WT males. However, *ex vivo* studies in females found no differences in the response to PE, ACh, or SNP in WT vs KO. Although KO females do have an increased response to KCl in the aorta, this alone seems unlikely to explain the blood pressure phenotype. However, female KOs manifest increased pulse wave velocity *in vivo*, indicating stiffer blood vessels. The fact that the female KO vessels appeared normal *ex vivo* but were stiffer *in vivo* implies that there may be a circulating factors or hormones required to manifest the phenotype.

### Human genetic analysis

In support of a role for this evolutionarily conserved OR in blood pressure regulation, OR51E1 was previously identified as a locus associated with DBP^44^. We report here that a (relatively) common OR51E1 variant (rs17224476; MAF=11.1%) is associated with increased DBP and affects blood pressure similarly in both men and women, whereas a rare OR51E1 variant (rs202113356; MAF=0.025%) effects blood pressure in the opposite direction in men as opposed to women. Notably, both the magnitude and the direction of the change in blood pressure for the rare variant is remarkably similar to the changes seen in male and female Olfr558 KO. It is also intriguing that the rare variant seems to have a stronger effect on increasing blood pressure in women <50 versus women >50. It is imperative to note that due to the rarity of this variant, the number of patients with this variant is quite small and thus we must interpret our conclusions appropriately according to the reduced statistical power. However, the presence of a variant in humans which differentially alters blood pressure by sex is supportive of the idea that our findings in mice are relevant to other species, including humans, especially when we observe such consistency and similarity in the effect estimates from our analyses in both humans and mice.

### Olfr558/OR51E1 trafficking and ligands

To explore how these two OR51E1 variants may alter the cell biology of OR51E1 signaling, we carried out *in vitro* studies. We previously published that the surface expression of the rare variant is reduced as compared to wild-type^46^, however, we find here that the common variant does not alter surface expression. We previously reported that butyrate is the best ligand for Olfr558/OR51E1, and that other ligands including IVA, VA, 3-MAV, 4-MAV, and CCA also activate Olfr558 and OR51E1^51^. In this study, we find that the activity of both the common and rare variants are reduced *in vitro* compared to wild-type OR51E1. Given that both variants reduce OR51E1 activity, yet the effects of the variants *in vivo* are different in each sex, it seems likely factors are missing from our *in vitro* system which prevent full recapitulation of OR51E1 biology. These factors may include sex chromosomal effects (HEK293T are XX), the absence of gonadal hormones, or downstream signaling partners. Of note, in our *in vivo* studies we were unable to detect any differences in Olfr558 expression (levels, or, localization) in males and females. Thus, there are likely other signals which “tell” the receptor whether it is in a male versus a female, and/or which alter its downstream signaling accordingly. Thus, moving forward, it is important to examine both a potential role for gonadal hormones to influence the receptor, and, whether there may be genes on the X or Y chromosome which alter some aspect of Olfr558 expression or signaling.

In conclusion, we have elucidated that an evolutionarily conserved OR is required for sex differences in blood pressure. In a mouse model, we find that Olfr558 KO females increase blood pressure, which is associated with increased arterial stiffness, whereas KO males decrease DBP due to alternations in renin as well as vascular reactivity. Uncovering the origin of sex differences in blood pressure regulation can help to move the field toward a more thoughtful approach to blood pressure management in men and women.

## List of abbreviations

Olfr558: olfactory receptor 558
WT: wildtype
KO: knockout
ORs: olfactory receptors
GPCRs: G-protein-coupled-receptors
OR51E1: olfactory receptor family 51 subfamily E member 1
SCFAs: short-chain fatty acids
PE: phenylephrine
SNP: sodium nitroprusside
ACh: acetylcholine
GFR: glomerular filtration rate
KW/BW: kidney weight/body weight
HW/BW: heart weight/body weight
OR51E1 A156T: OR51E1 variant rs2021113356
OR51E1 S10N: OR51E1 variant rs17224476
PFA: paraformaldehyde
HEK 293T: human embryonic kidney 293T
RTP1S: receptor transporting protein 1S
Golf: olfactory G protein
IVA: isovaleric acid
VA: valeric acid
3-MVA: 3-methylvaleric acid
4-MVA: 4-methylvaleric acid
CCA: cyclobutene-carboxylic acid

## Acknowledgements

We would like to thank Jason Sanchez, Dylan Sarver, Jie Sun, Shuai Wu, Brian Poll, and P. Richard Grimm for their advice and help. We also thank Pluznick Lab members, including Nathan Zaidman, Muhammad Umar Cheema, Mackenzie Kui, Brittni Moore, Brian Poll, Jason Sanchez, and Tilmira Smith for helpful discussion, suggestions, and comments. This research has been conducted using the UK Biobank Resource under application no. 236.

## Funding

This work was supported by R56DK107726 (to JL Pluznick). H.R. Warren acknowledges the funding of the NIHR Cardiovascular Biomedical Centre at Barts and The London, Queen Mary University of London.

## Ethics statement

The UK Biobank study has approval from the North West Multi-Centre Research Ethics Committee. Any participants from UK Biobank who withdrew consent were removed from our analysis.

## Disclosures

The authors have no competing interests to declare.

## Authors’ Contributions

J.L.P., J.X. and L.S. conceived and designed the research. J.X., R.C., H.W., and K.G. performed the experiments. J.X., R.C., and H.W. analyzed the data. J.L.P., J.X., and L.S. interpreted the data. J.X. and J.L.P. wrote the manuscript. J.L.P. provided funding. All authors read and approved the final manuscript.

## Notes

### Competing Interest Statement

The authors have declared no competing interest.

