## Supplemental Figures and Tables for "An evolutionarily conserved olfactory receptor is required for sex differences in blood pressure"

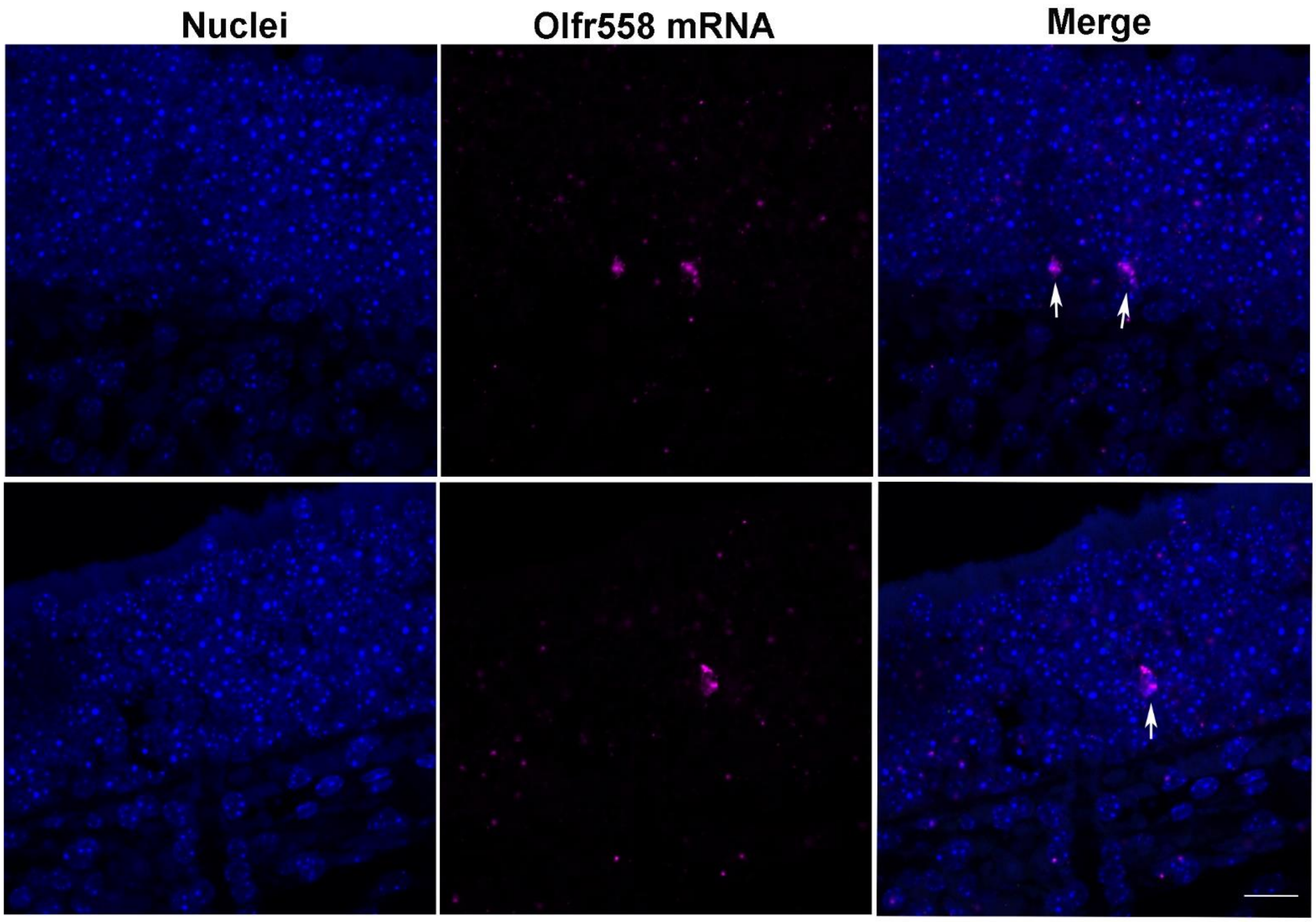


**Figure S1. Olfr558 expression in the OE.** Olfr558 mRNA staining (purple, white arrow) is observed in zone 1 of the olfactory epithelium. Nuclear stain is blue; two different fields of view are shown. Scale bar: 20 μm.

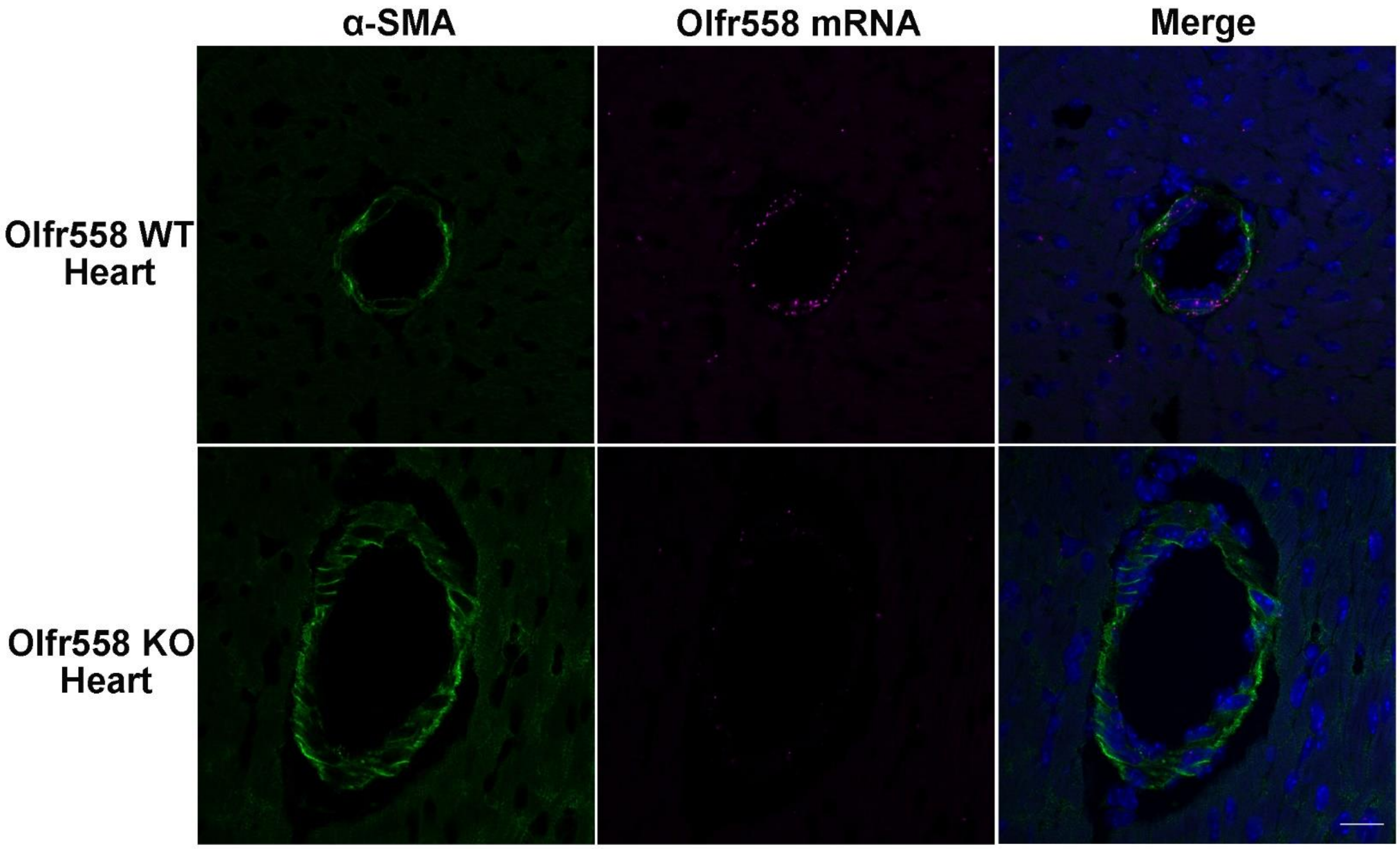


**Figure S2. Olfr558 expression in vascular smooth muscle cells in the heart.** RNAScope and immunostaining demonstrate that Olfr558 mRNA (purple) is expressed in blood vessels labeled by α-SMA (green, blood vessel marker) in Olfr558 WT but not KO heart. Nuclear stain is blue. Scale bar: 20 μm.

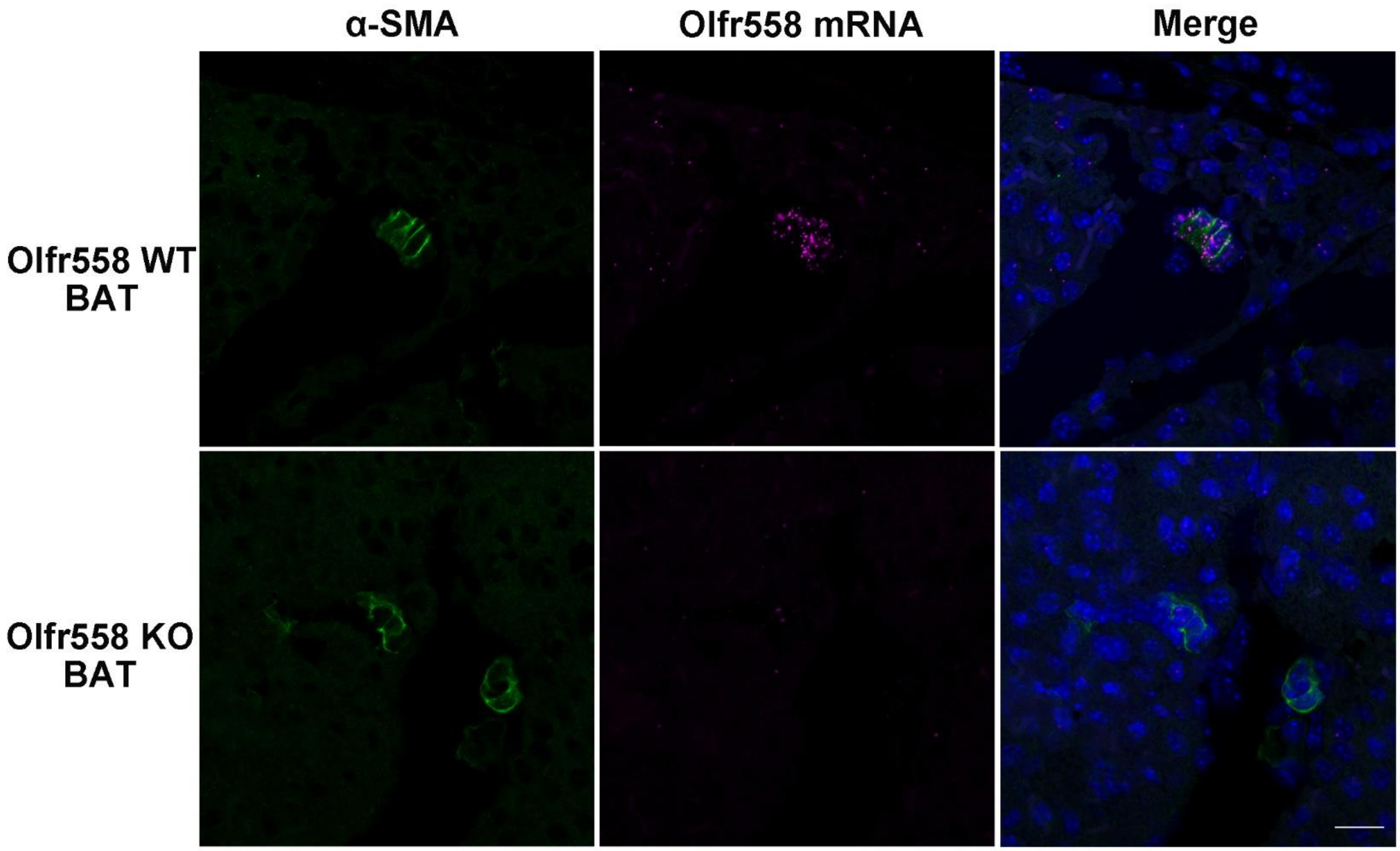


**Figure S3. Olfr558 expression in vascular smooth muscle cells in brown adipose tissue (BAT).** RNAScope and immunostaining demonstrate that Olfr558 mRNA (purple) is expressed in blood vessels which are labeled by α-SMA (green, blood vessel marker) in BAT from WT but not KO mice. Nuclear stain is blue. Scale bar: 20 μm.

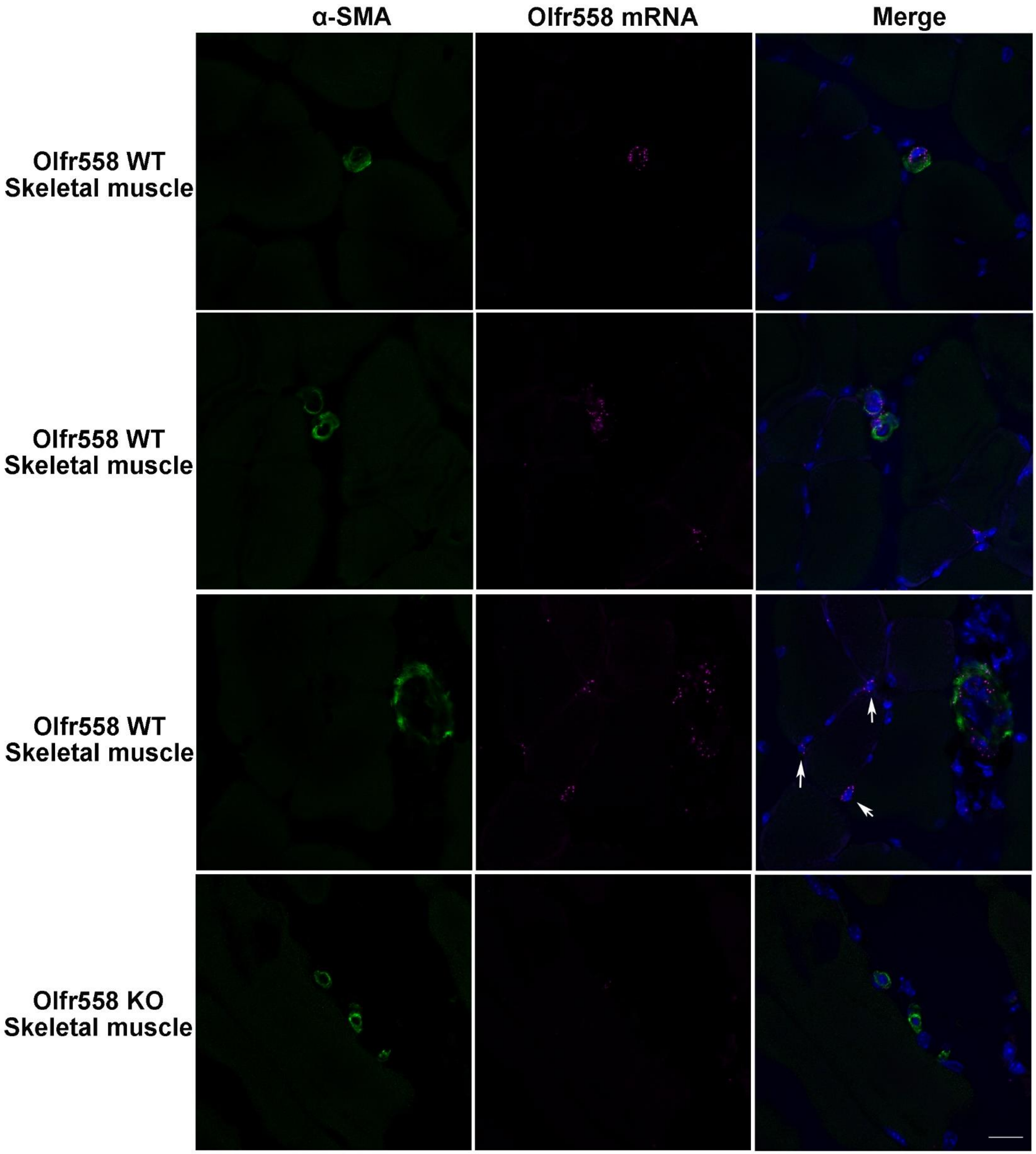


**Figure S4. Olfr558 is primarily expressed in blood vessels in skeletal muscle.** RNAScope and immunostaining demonstrate that Olfr558 mRNA (purple) is expressed in blood vessels which are labeled by α-SMA (green, blood vessel marker) in Olfr558 WT skeletal muscle, as well as a minority cell type which is negative for α-SMA (white arrows). Olfr558 mRNA staining is absent in Olfr558 KO. Nuclear stain is blue. Scale bar: 20 μm.

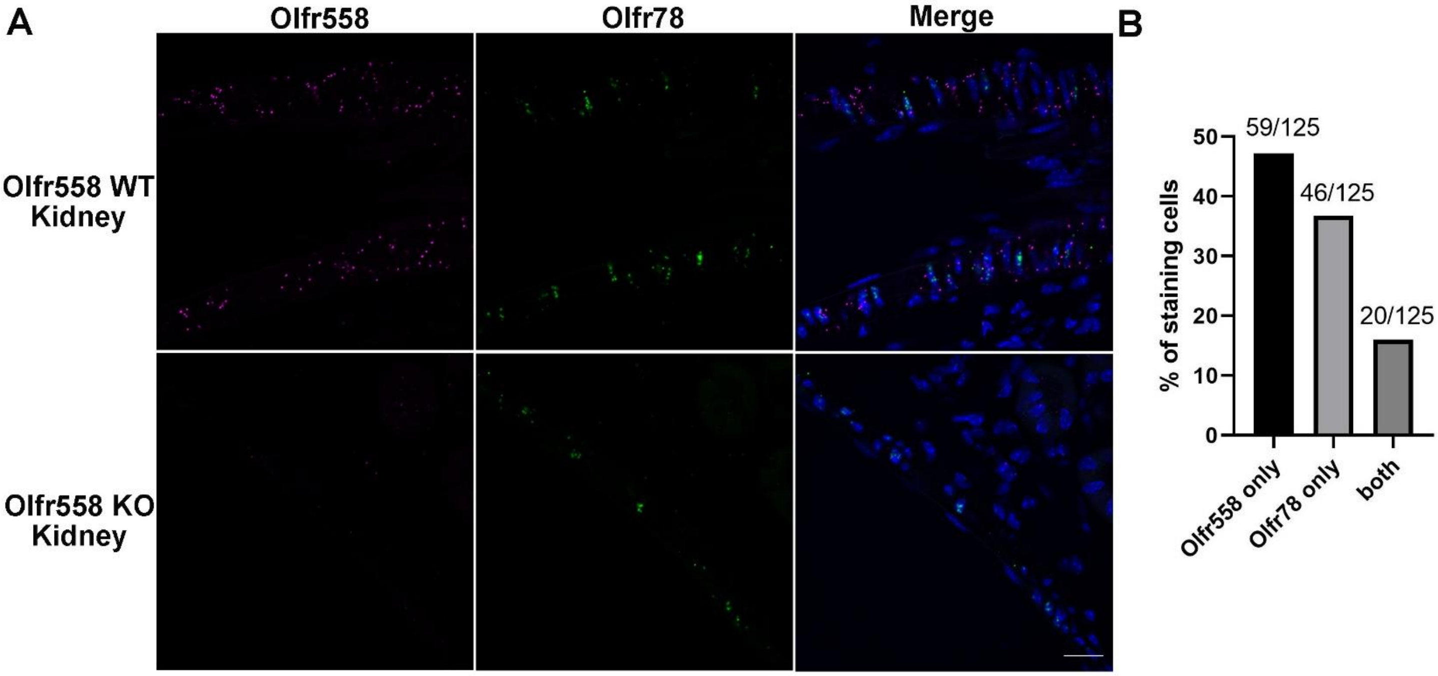


**Figure S5. Olfr558 and Olfr78 expression in the kidney. (A)** Olfr558 (purple) and Olfr78 mRNA staining (green) are both observed in a blood vessel from Olfr558 WT kidney, but Olfr558 mRNA staining is absent in Olfr558 KO kidney. Nuclear stain is blue. **(B)** Quantification of Olfr558 and/or Olfr78 staining cells in Olfr558 WT renal blood vessels. Quantification performed on n=125 vascular smooth muscle cells from n=8 WT kidney sections. Scale bar: 20 µm.

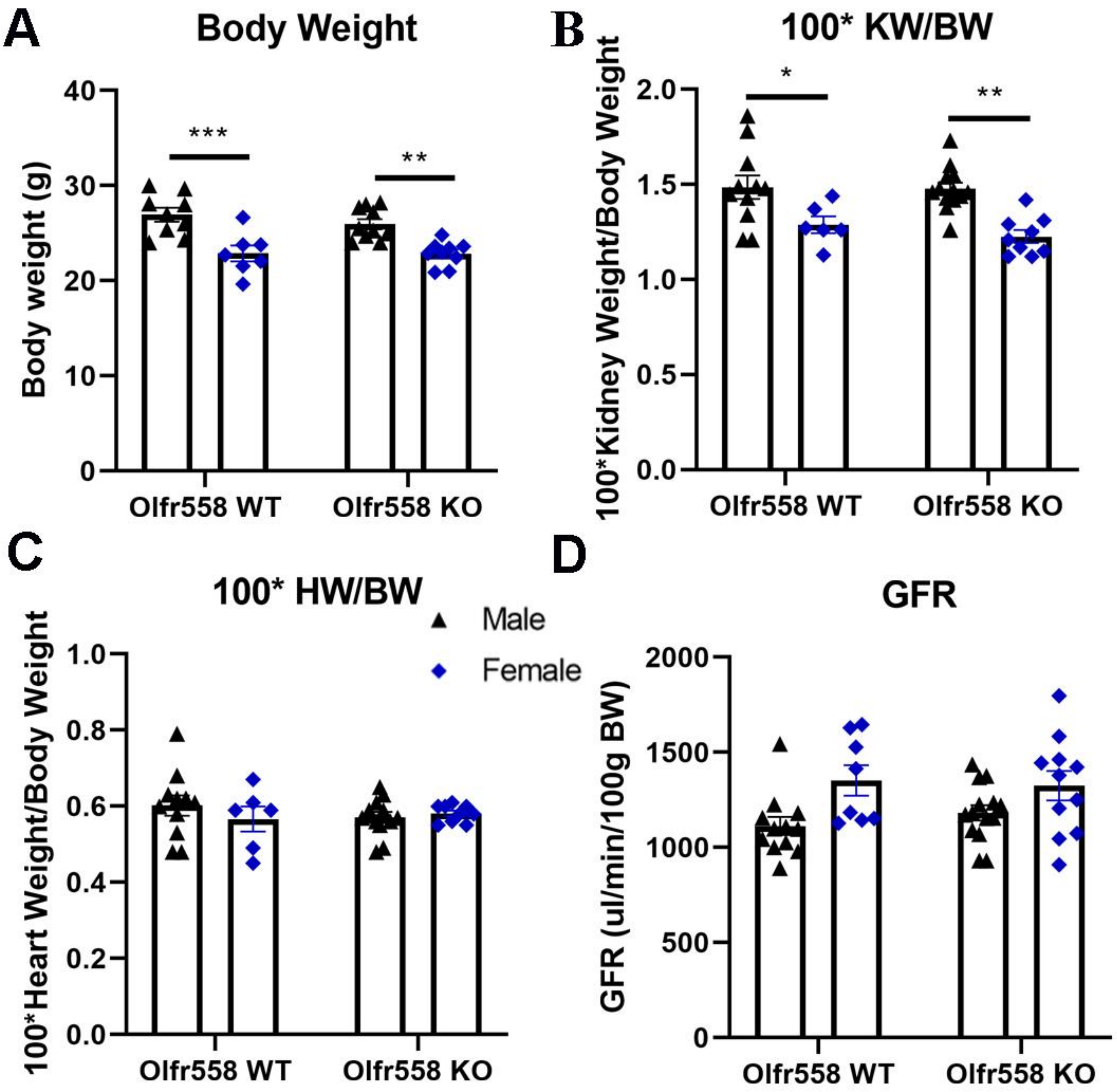


**Figure S6. Body weight, kidney weight/body weight (KW/BW), heart weight/body weight (HW/BW), and GFR in Olfr558 mice. (A)** Both WT and KO mice have sex differences in body weight; however, there are no genotypic differences. **(B)** Similarly, there are sex differences but not genotypic differences for KW/BW. **(C)** HW/BW shows no significant difference between genotypes or sexes. **(D)** Glomerular filtration rate (GFR) indicates no differences between genotypes or sexes. *$p<0.05$, **$p<0.01$; ***$p<0.001$, ****$p<0.0001$ by genotype (Olfr558 WT vs. KO) and sex (Male vs. Female) using two-way ANOVA. ns: non-significant. n=9-13 for males, n=7-11 for females, mice were 10-12 weeks of age.

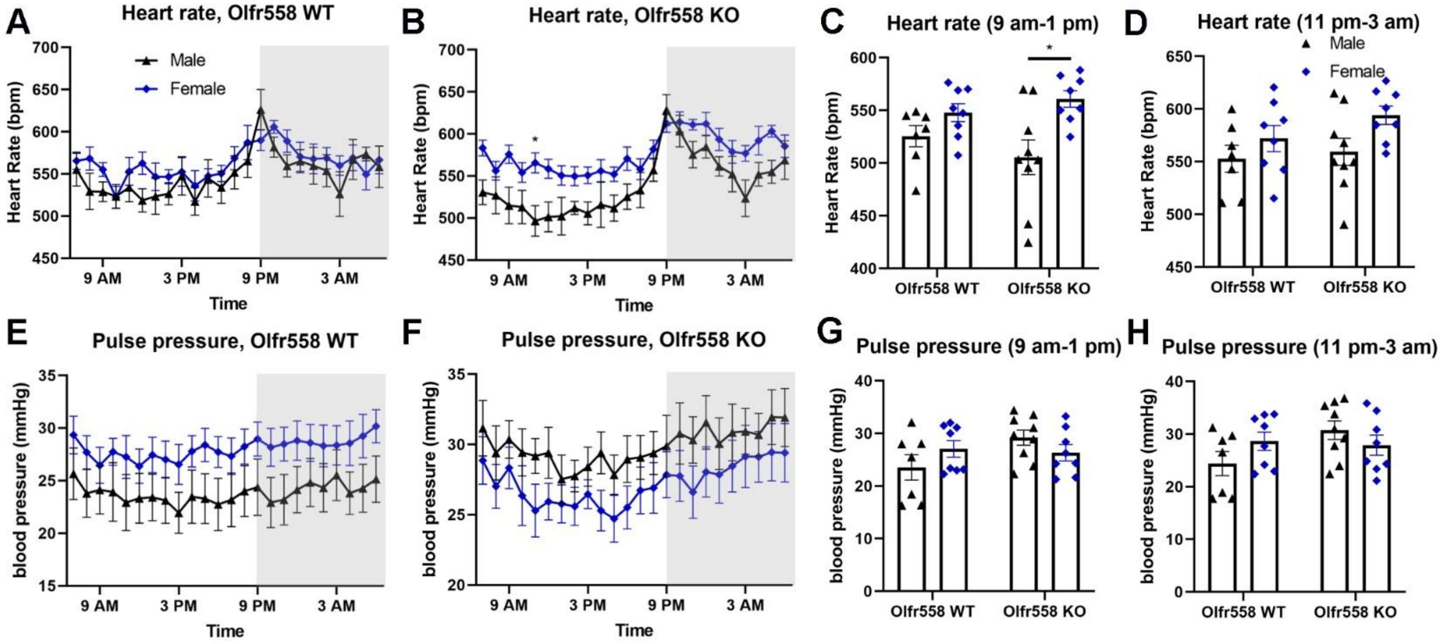


**Figure S7. Olfr558 regulation in heart rate and pulse pressure.** Telemetry was utilized to measure 24 h heart rate and blood pressure. Heart rate **(A, B)** and pulse pressure **(E, F)** averages are from 14 h light and 10 h dark cycles. Grey areas depict the activity (dark) phase and white areas depict the resting (light) phase. Light phase: 7 am-9 pm; dark phase: 9 pm-7am. Heart rate data (**C, D**) and pulse pressure (**G, H**) data are also shown as averages from 9 am-1 pm (light cycle) and 11 pm-3 am (dark cycle). Male WT: n=7, male KO: n=9. Female WT and KO: n=8. *$p<0.05$ by sex (Male vs. Female) using two-way ANOVA.

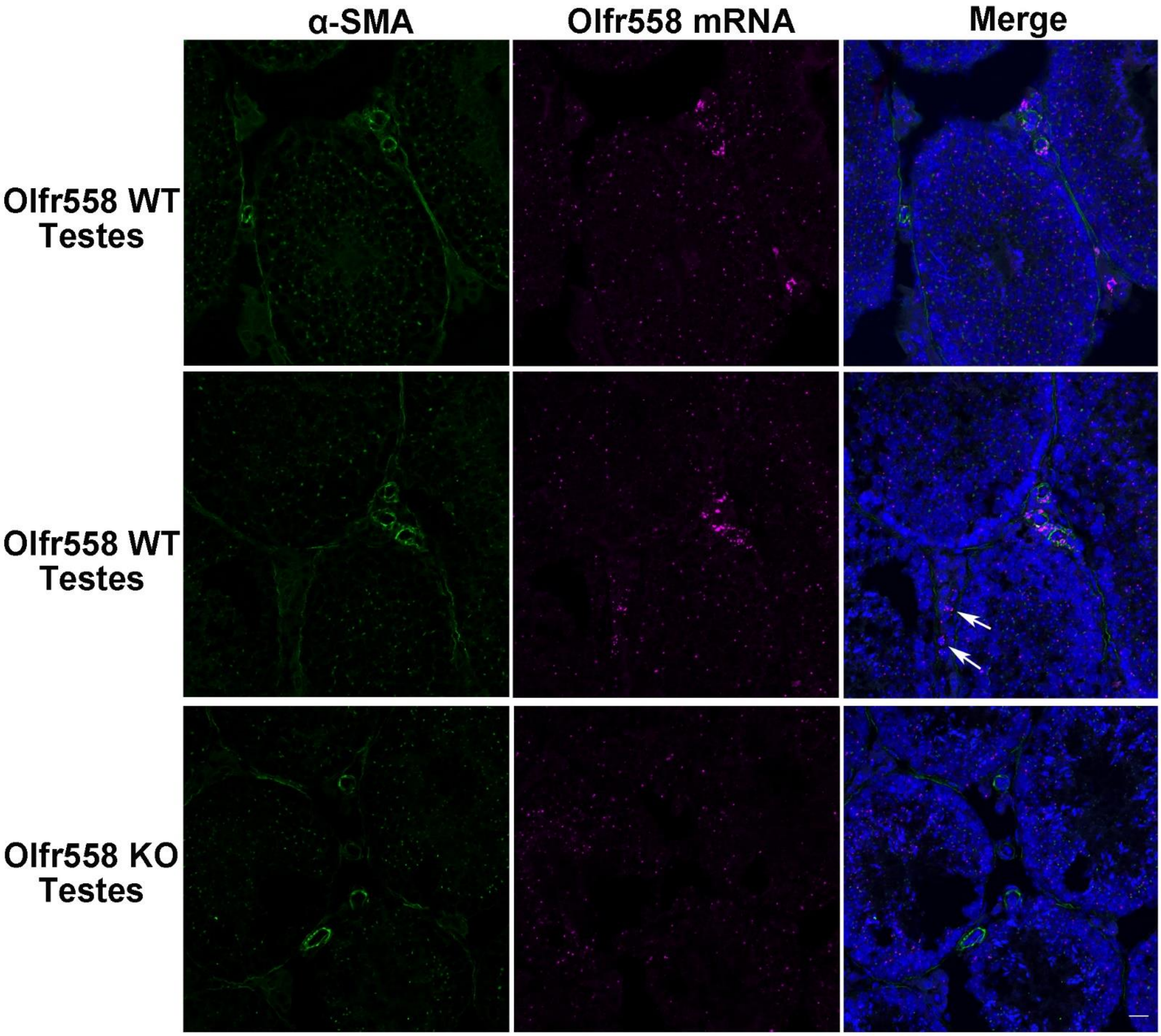


**Figure S8. Olfr558 is primarily expressed in blood vessels in testes.** RNAScope and immunostaining demonstrate that Olfr558 mRNA (purple) is expressed in blood vessels which are labeled by α-SMA (green, blood vessel marker) in Olfr558 WT testes, as well a minority cell type which is negative for α-SMA (white arrow). Olfr558 mRNA staining is absent in Olfr558 KO. Nuclear stain is blue. Scale bar: 20 μm.

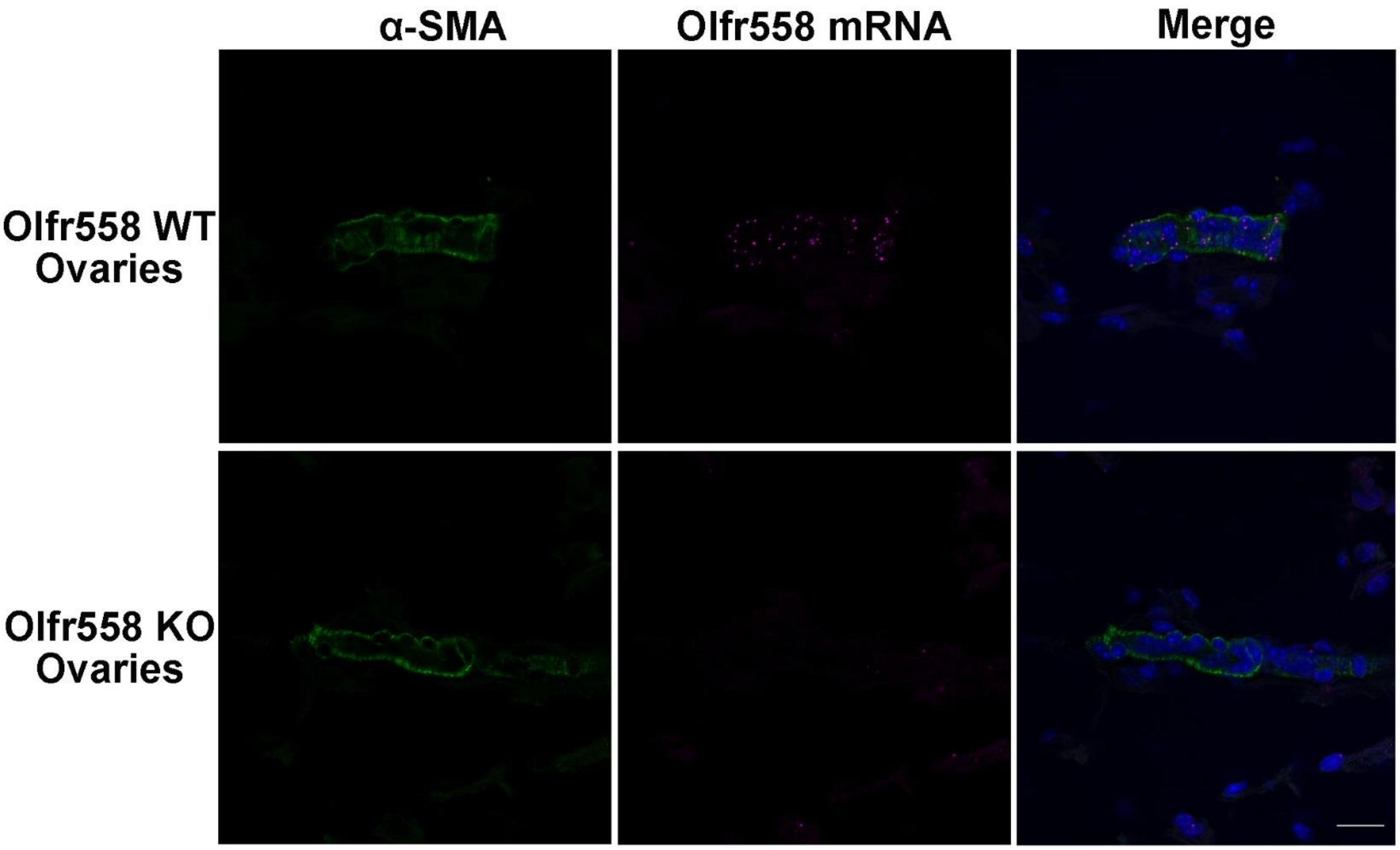


**Figure S9. Olfr558 is expressed in blood vessels in ovaries.** RNAScope and immunostaining demonstrate that Olfr558 mRNA (purple) is expressed in blood vessels which are labeled by α-SMA (green, blood vessel marker) in WT but not KO ovaries. Nuclear stain is blue. Scale bar: 20 μm.

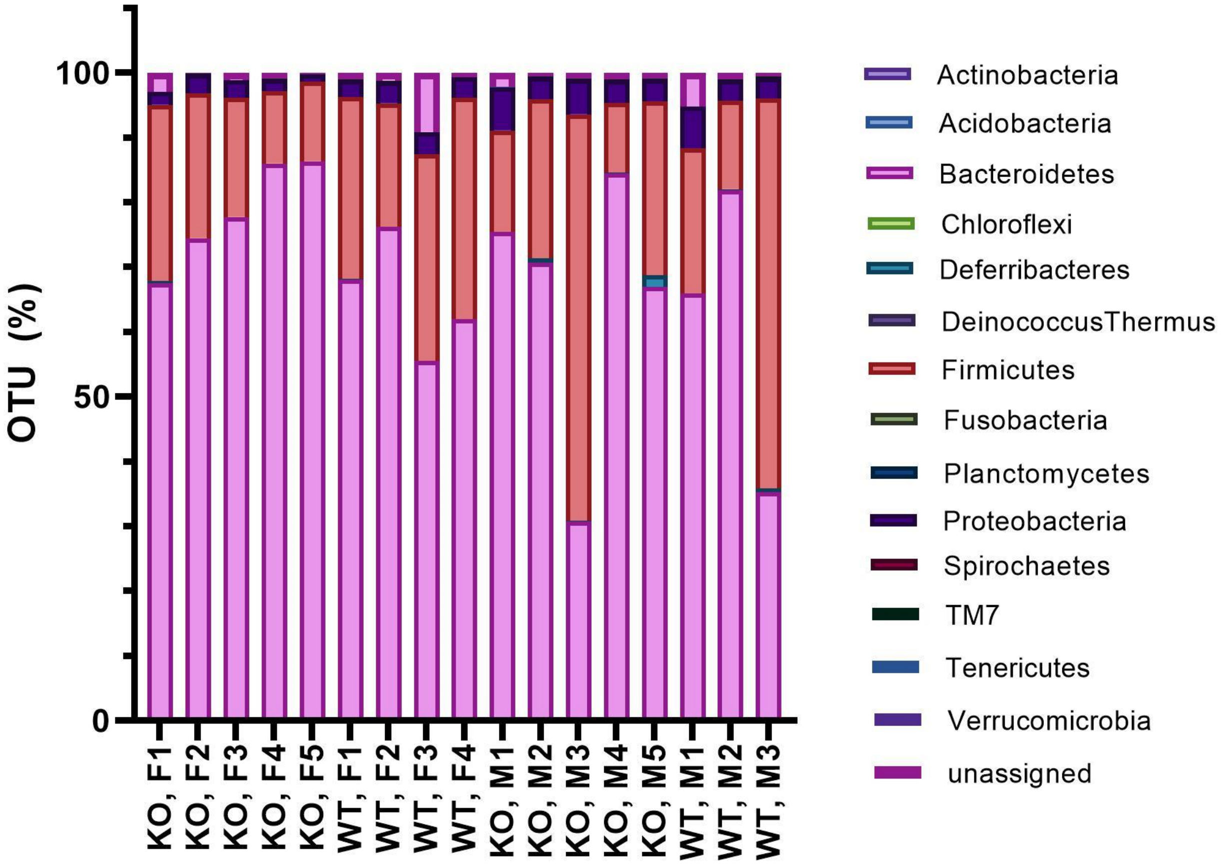


**Figure S10**. **Microbial sequencing (16s) from feces of Olfr558 WT and KO mice (both males and females)**. Analysis on the phylum level revealed no differences between Olfr558 WT and KO. However, proteobacteria is higher in male KO versus female KO (*p*=0.03 by one-way ANOVA). WT: Olfr558 WT, KO: Olfr558 KO, F: female, M: male. n=5 female KO, n=4 female WT, n=5 male KO, n=4 male KO.

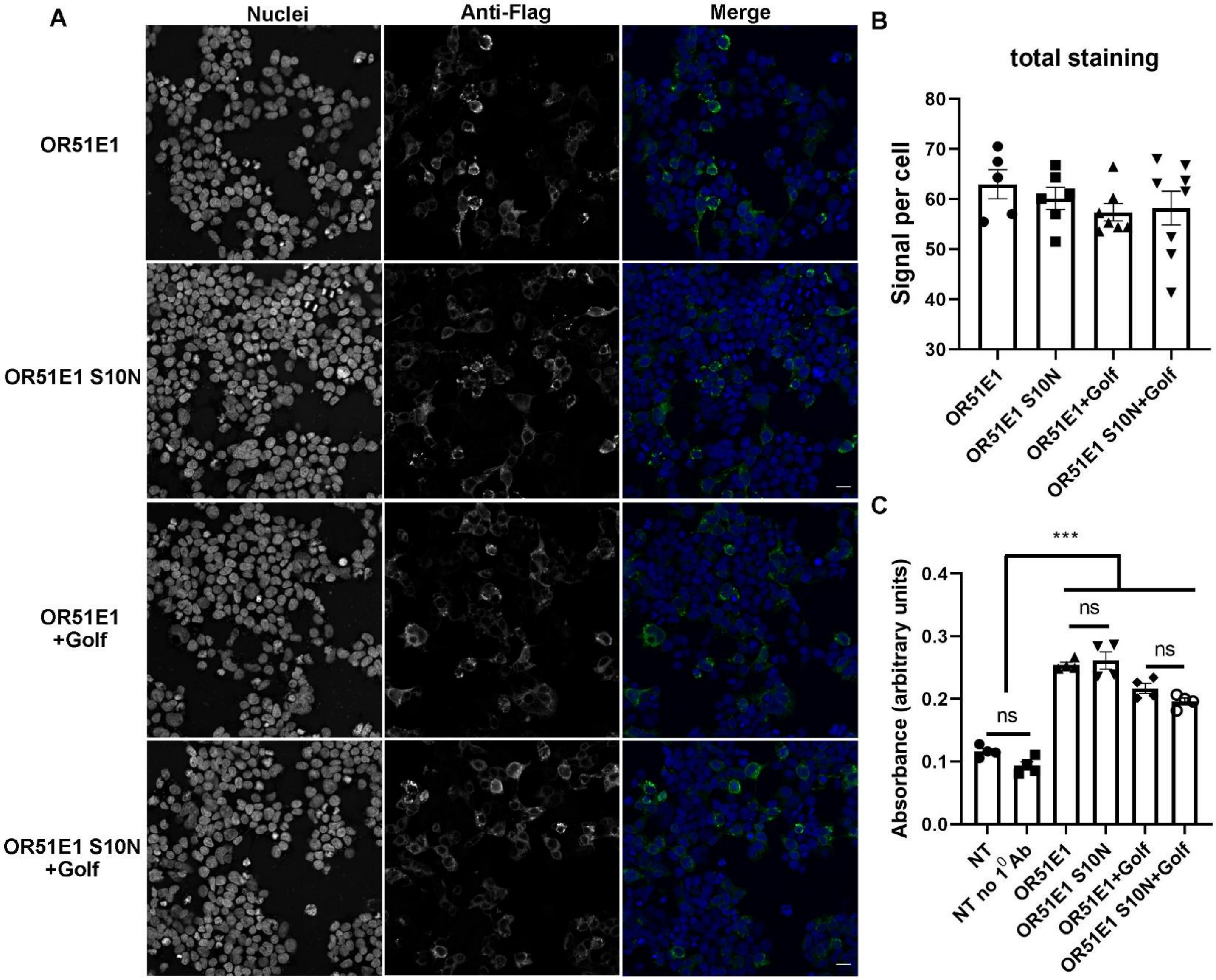


**Figure S11. OR51E1 S10N does not alter total protein expression.** Representative images **(A)** and quantification of total protein expression are shown **(B)**. HEK 293T cells, transfected with OR51E1 or OR51E1 S10N (with or without Golf), are stained with mouse anti-Flag antibody (permeabilized total staining, green in merge). Scale bar: 20 μm. Quantification of signal per cell for OR total protein expression (**B**) is shown where each data point represents the average signal per cell for a given field of view. There is no significant difference between OR51E1(+/-Golf) vs. OR51E1 S10N (+/-Golf) by one-way ANOVA. Total OR protein expression is also detected using anti-Flag antibody in HEK 293T cells transfected with OR51E1 or OR51E1 S10N (with or without Golf) by an ELISA assay **(C)**. NT (non-transfected cells) and NT no 1$^{o}$ Ab (non-transfected cells without primary antibody) are used as controls. These data also show that OR51E1 S10N does not affect the total protein expression. ***$p$<0.001 vs. OR by one-way ANOVA.

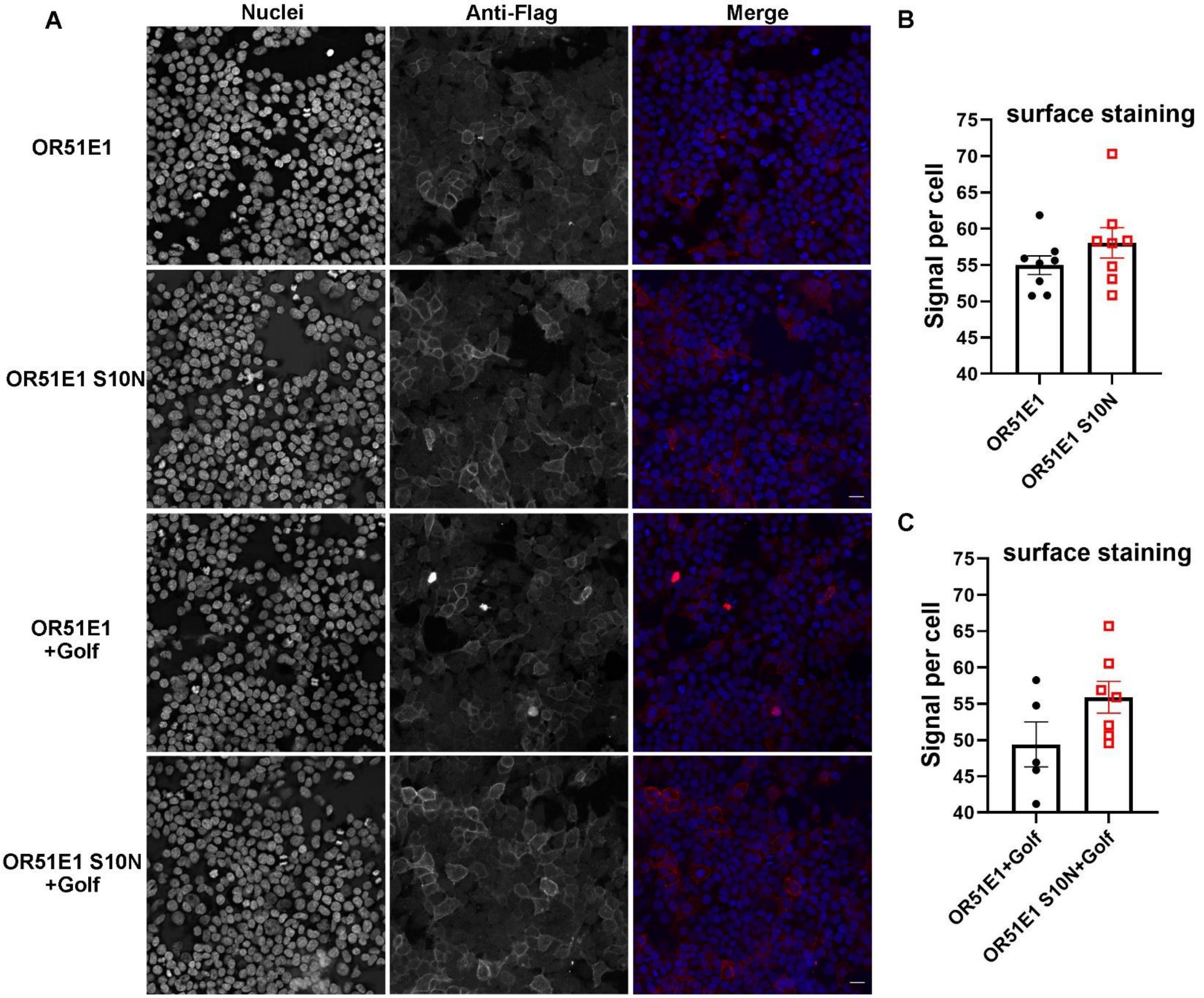


**Figure S12. OR51E1 S10N does not alter cell surface expression**. Representative images (**A**) and quantification of cell surface expression (**B-C**) are shown. HEK 293T cells, transfected with OR51E1 or OR51E1 S10N (with or without Golf), are stained with rabbit anti-Flag antibody (unpermeabilized surface staining, red in merge). The flag tag is on the N-terminus, and thus the flag is extracellular (and able to be labeled) only if the receptor reaches the cell surface). Scale bar: 20 μm. Quantification of signal per cell for OR surface expression (**B-C**) is shown where each data point represents the average signal per cell for a given field of view. Non-significant difference between OR51E1(+Golf) vs. OR51E1 S10N (+Golf) by t-test. n=3 independent experiments. Data are shown as mean ± SEM.

### Supplementary Tables

**Supplementary Table 1. Plasma electrolytes in Olfr558 WT and KO mice.**

| | M, WT | M, KO | F, WT | F, KO | ***p*** **(male: WT vs KO)** | ***p*** **(female: WT vs KO)** | ***p*** **(all, WT vs. KO)** |
|---|---|---|---|---|---|---|---|
| Na, mM/L | 141.6±0.26 | 141.5±0.30 | 142.2±0.11 | 142.0±0.16 | 0.97 | 0.80 | 0.79 |
| Cl, mM/L | 116.1±0.35 | 113.7±0.43 | 114.3±0.18 | 113.4±0.25 | 0.19 | 0.45 | 0.15 |
| iCa, mM/L | 1.07±0.01 | 1.08±0.02 | 1.09±0.01 | 1.11±0.01 | 0.83 | 0.73 | 0.75 |
| $TCO_2$, mM/L | 18.3±0.29 | 19.5±0.30 | 18.8±0.14 | 20.3±0.19 | 0.38 | 0.10 | 0.09 |
| Glucose, mg/dl | 223.4±3.39 | 241.3±3.14 | 219.8±2.28 | 216.9±3.42 | 0.25 | 0.85 | 0.46 |
| BUN, mg/dl | 21.4±0.57 | 21.7±0.43 | 19.4±0.38 | 21.6±0.67 | 0.89 | 0.41 | 0.41 |
| Creatinine, mg/dl | <0.2 | <0.2 | <0.2 | <0.2 | | | |
| Hematocrit, %PCV | 40.9±0.29 | 39.7±0.33 | 41.3±0.18 | 39.8±0.21 | 0.40 | 0.19 | 0.11 |
| Hemoglobin, g/dl | 13.9±0.10 | 13.5±0.11 | 14.7±0.17 | 13.5±0.07 | 0.42 | 0.22 | 0.13 |
| # of mice | n=11 | n=10 | n=17 | n=10 | | | |

Blood chemistries of Olfr558 WT and KO mice are similar. M: male, F: female; iCa: ionized calcium; $TCO_2$: total carbon dioxide; BUN: blood urea nitrogen; PCV: packed cell volume. ANOVA analysis for the male WT, male KO, female WT, female KO. A t-test for all WT vs KO analysis.

**Supplementary Table 2. Demographic analysis of data from UK Biobank** (attached excel file)**.**

**Supplementary Table 3. Association Results for Analysis of Diastolic Blood Pressure in UK Biobank. (A) for rs17224476 lead variant; (B) for rs202113356 rare variant.** Genotype model analysis compares GG non-carriers vs AG/AA carriers of the minor allele. "BETA": effect estimate in mmHg for the effect of the carrier AG/AA group; "P" = P-value (in bold if P<0.5); "SE" = Standard Error of Beta; "CI" = 95% Confidence Interval of effect; "N" = Sample sizes. (attached excel file)
